# The Structured RNA-binding Domains and Condensation Capacity of FUS Shape its RNA-binding Landscape and Function

**DOI:** 10.64898/2026.01.12.699057

**Authors:** Daniel Jutzi, Juan Alcalde, Saskia Hutten, Fatmanur Tiryaki, Benjamin Davies, Helene Plun-Favreau, Christopher R. Sibley, Dorothee Dormann, Marc-David Ruepp

## Abstract

RNA-binding proteins (RBPs) are key regulators of gene expression and often contain intrinsically disordered regions that drive biomolecular condensation. Yet, how condensation affects RBP specificity and function remains poorly defined. Here, we use strategically designed point mutations to selectively impair canonical RNA-binding and condensation of the amyotrophic lateral sclerosis (ALS)-linked RBP Fused in Sarcoma (FUS). Using automated high-content imaging, we show that both properties shape nuclear ribonucleoprotein condensates and govern distinct aspects of FUS function in the DNA damage response. Transcriptome-wide mapping of FUS-RNA interactions reveals that the canonical RNA-binding domains recognise G-rich and C-rich motifs, whereas condensation selectively enhances binding to G-rich and structured sequences, often acting in concert with canonical RNA-binding to control transcriptional programs and splicing decisions. These findings provide a mechanistic framework for how FUS integrates condensation with sequence-specific RNA recognition to orchestrate nuclear organisation, genome stability and RNA metabolism. Furthermore, our rigorously validated mutants offer a platform for future mechanistic dissection of ALS pathogenesis and the development of targeted therapeutic strategies.

**Highlights:**

- FUS condensation and RNA-binding can be functionally uncoupled with targeted point mutations
- Condensation drives FUS recruitment and assembly of nuclear RNP condensates
- FUS condensation and RNA-binding play distinct roles in the DNA damage response
- The structured RNA-binding domains confer specificity to G-rich and C-rich motifs
- Condensation enhances FUS binding to G-rich, structured RNA elements
- FUS condensation and canonical RNA-binding synergistically regulate gene expression

## Introduction

RNA-binding proteins (RBPs) are key regulators of gene expression^1–3^. Besides structured RNA-binding domains, they frequently contain intrinsically disordered regions (IDRs) that engage in multivalent, low-specificity interactions that drive biomolecular condensation, a dynamic process that operates across scales, giving rise to non-stochiometric molecular assemblies as well as micron-scale membraneless compartments^4–7^. Yet, our understanding of how condensation interfaces with sequence-specific RNA recognition and shapes the physiological function of RBPs remains remarkably limited.

Fused in Sarcoma (FUS) is a ubiquitously expressed RBP that is co-transcriptionally recruited to pre-mRNA and accompanies transcripts throughout their life cycle, regulating key processes such as transcription, alternative splicing, polyadenylation and mRNA transport^8–12^. Besides its role in RNA metabolism, FUS is also recruited to sites of DNA damage and contributes to multiple DNA repair pathways^13–17^. Mutations in the FUS gene cause a rare form of ALS, often leading to an aggressive disease phenotype with juvenile onset^18,19^. The most common mutations disrupt the nuclear localisation signal (NLS) and thus result in FUS mislocalisation to the cytoplasm^20^ where the protein is thought to exert a toxic gain-of-function through mechanisms that remain poorly understood^21,22^, hindering the development of targeted therapies.

FUS contains two structured RNA-binding domains: an RNA recognition motif (RRM) and a zinc finger (ZnF) domain. *In vitro*, using purified components, these domains preferentially recognise a bi-partite motif consisting of a YNY motif-containing stem loop - bound by the RRM - and a neighbouring GGU trinucleotide, which is contacted by the ZnF domain^23^. In cells, however, FUS binding is not limited to these specific motifs but instead extends to a broad range of sequences which are widely scattered along nascent transcripts^10,11^. This suggests that FUS-RNA interactions are not solely guided by its canonical RNA-binding domains. Indeed, some studies implicate the intrinsically disordered arginine-glycine-glycine (RGG) domains of FUS as major contributors to RNA-binding^24–26^. This raises important questions: To what extent do the canonical RNA-binding domains contribute to FUS-RNA interactions *in vivo*, how do they shape its sequence specificity, and how do these interactions influence the function of FUS in regulating gene expression?

Another key functional property of FUS is its propensity for condensation. Above a critical saturation concentration, purified FUS undergoes phase separation^27^, a process where a homogeneous solution of molecules spontaneously segregates into two coexisting phases: a dense phase enriched for these molecules and a dilute phase depleted of them. Phase separation is driven by multivalent interactions throughout the intrinsically disordered regions, with a prominent role played by tyrosine residues in the N-terminal low-complexity (LC) region and arginine residues in the C-terminal RGG repeats^28–32^. Additional elements which have been linked to FUS condensation are the N-terminal ‘core’ region (amino acids 39 – 95)^33^ and the reversible amyloid cores ^37^ SYSGYS^42^ (RAC1) and ^54^ SYSSYG^59^ (RAC2)^34^, which mediate the reversible formation of β-sheet rich fibrils. In cells, FUS partitions into membrane-less condensates that are thought to arise through phase separation of their constituents^35^. These include ribonucleoprotein (RNP) granules such as paraspeckles, which regulate nuclear RNA retention^36^, and nuclear gems/Cajal bodies, which promote maturation and assembly of small nuclear ribonucleoproteins (snRNPs)^37^ - both of which rely on FUS for their formation^38–41^. However, the functional implications of FUS condensation remain poorly understood. Key outstanding questions include: How does FUS condensation influence the composition and assembly of nuclear ribonucleoprotein (RNP) granules? Does condensation modulate the RNA-binding landscape of FUS? And to what extent is condensation required for the functions of FUS in gene regulation and the DNA-damage response?

In this study, we address these key questions using strategically designed FUS point mutations that selectively impair either condensation or canonical RNA-binding. We demonstrate that both properties are essential for the formation of Cajal bodies, whereas condensation specifically enhances the recruitment of FUS to PSPC1-positive paraspeckles and perinucleolar foci. Following DNA damage, FUS condensation facilitates its rapid accumulation at DNA damage sites and promotes the subsequent recruitment of histone deacetylase 1 (HDAC1), a chromatin-modifying enzyme that promotes non-homologous end joining (NHEJ)^42^. Under conditions of severe genotoxic stress, condensation mediates extensive FUS translocation to the nucleolus. In contrast, canonical RNA-binding by FUS is essential for maintaining the expression of key NHEJ regulators, thereby safeguarding effective double-strand break (DSB) repair. Using streamlined individual-nucleotide resolution cross-linking and immunoprecipitation (siCLIP), we find that condensation and canonical RNA-binding modulate binding to distinct subsets of FUS binding sites and identify features that confer this sensitivity: G-rich and C-rich motifs promote binding via the canonical RNA-binding domains, whereas condensation selectively strengthens binding to G-rich and structured RNA sequences but is dispensable for binding to C-rich motifs. Finally, our transcriptomic data highlight a cooperative interplay between canonical RNA-binding and condensation in orchestrating FUS-dependent gene expression and alternative splicing.

## Results

### Identification of RNA-binding and condensation-deficient FUS mutants

To dissect how canonical RNA-binding and condensation contribute to FUS function, we first sought to identify and validate point mutations that selectively impair either process. We focused on mouse FUS to ensure compatibility and physiological relevance with *in vivo* systems and provide a foundation for future studies in animal models. Given that the RRM and zinc finger domains are identical between human and mouse, the canonical RNA-binding deficient mutant (hereafter referred to as ‘RBdef FUS’) was designed based on our previously characterised human construct (Figures 1A and 1B). This construct harbours seven point mutations, informed by NMR structural insights, that preserve domain folding but markedly reduce binding to diverse RNA sequences *in vitro* using purified components^23,43^.

**Figure 1.**
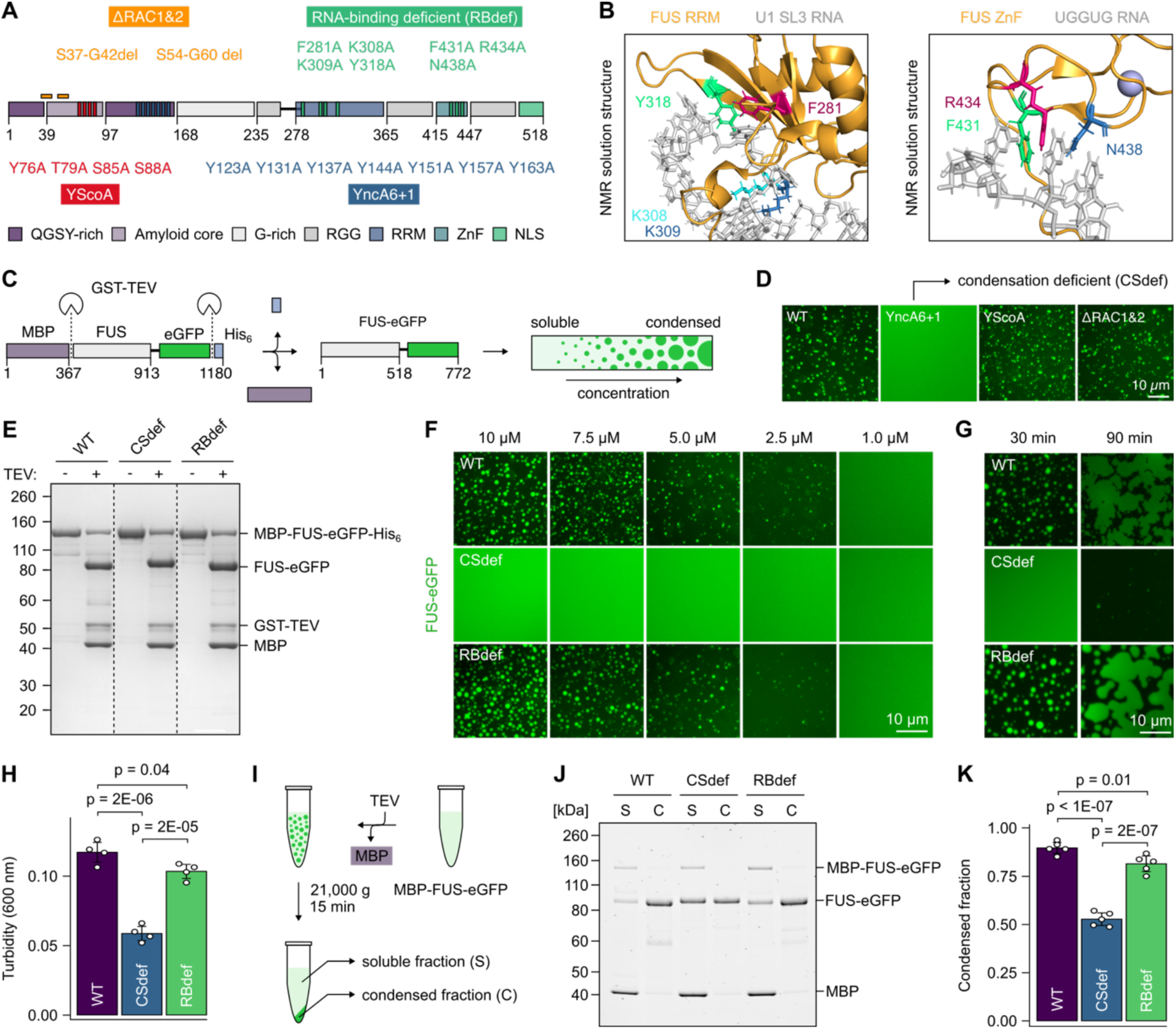
Identification of RNA-binding and condensation-deficient FUS mutants. (A) Schematic representation of candidate FUS mutants identified from literature. (B) NMR solution structures of the FUS RNA recognition motif (RRM, PDB: 6SNJ) and zinc finger domain (ZnF, PDB: 6G99) bound to RNA (grey), highlighting the seven residues (coloured) mutated to disrupt canonical RNA binding. (C) Schematic of the fluorescent droplet assay. Phase separation is induced by TEV-mediated cleavage of the N-terminal maltose-binding protein (MBP) solubility tag. (D) Fluorescence microscopy images of droplet formation at 5 µM protein concentration, 30 min after TEV cleavage. Scale bar, 10 µm. (E) Coomassie-stained SDS-PAGE gel confirming efficient cleavage of MBP-FUS-eGFP-His₆ constructs 30 min after addition of GST-tagged TEV protease. (F) Droplet formation visualized at varying FUS-eGFP concentrations, 30 min after TEV cleavage. Scale bar, 10 µm. (G) Time course of droplet formation at 7.5 µM FUS-eGFP, imaged 30 and 90 min after TEV cleavage. Scale bar, 10 µm. (H) Turbidity at 600 nm used to quantify phase separation of 7 µM FUS-eGFP, 30 min after TEV cleavage. Data represent mean ± SD of N = 4 independent biological replicates. Statistical significance was determined by one-way ANOVA followed by Tukey’s Honest Significant Difference (HSD) test for multiple comparisons. (I) Schematic of the sedimentation assay. Following TEV cleavage, condensed and soluble protein fractions were separated by centrifugation. (J) SyproRuby-stained SDS-PAGE gel showing partitioning of FUS-eGFP between soluble (S) and condensed (C) fractions. 2 µM protein, 60 min after cleavage with His_6_-tagged TEV. (K) Quantification of the condensed fraction (condensed / total) from panel (J). Data represent mean ± SD of N = 5 independent biological replicates. Statistical significance was determined by one-way ANOVA followed by Tukey’s HSD test.

Sequence differences between human and mouse FUS, as well as the ALS-associated NLS mutation P517L (P525L in humans), did not affect phase separation of recombinant full-length FUS *in vitro* (Figure S1A). To probe a condensation-deficient mutant, we therefore engineered a panel of mouse FUS P517L variants targeting key determinants of condensation and self-assembly (Figure 1A), guided by published structural and biophysical insights: FUS with wild-type (WT) low complexity domain, a mutant with tyrosine substitutions downstream of the fibril core region (YncA6+1)^29^, a variant lacking the reversible amyloid core motifs (ΔRAC1&2)^34^, and a mutant with destabilised fibril core folding (YScoA)^33^. Following expression and purification of MBP-FUS-eGFP-His₆ fusion proteins from *E. coli*, phase separation was induced by TEV-mediated cleavage of the MBP solubility tag (Figure 1C). We selected the eGFP-tag because it only minimally affects FUS condensation across a wide range of pH and salt concentrations^44^. Among the tested constructs, only the YncA6+1 mutant (hereafter referred to as ‘CSdef FUS’) remained fully soluble under conditions where WT FUS robustly phase separated (Figure 1D; Figure S1B).

To further characterise CSdef FUS and confirm that the canonical RNA-binding mutations do not intrinsically affect phase separation, we compared WT, CSdef, and RBdef FUS side by side. After 30 minutes of TEV cleavage, both WT and RBdef FUS exhibited robust, concentration-dependent droplet formation, initiating at ∼2.5 µM, consistent with previous reports^31^ (Figures 1E and 1F; Figure S1C). In stark contrast, CSdef FUS failed to form droplets at any concentration tested, indicating that its saturation concentration (C_sat_) is elevated by at least 4-fold relative to WT. After 90 minutes, WT and RBdef droplets matured through fusion and wetting on the glass surface, whereas CSdef FUS displayed only sparse, small droplets, underscoring its severely delayed and impaired phase separation (Figure 1G). The condensation defect of CSdef FUS was independently validated by turbidity measurements (Figure 1H; Figure S1D) and sedimentation assays (Figures 1I, 1J and 1K), both demonstrating that the tyrosine substitutions markedly reduce the fraction of FUS in the condensed state (∼2-fold in turbidity assays, ∼1.7-fold in sedimentation assays).

We next examined how RNA influences phase separation by performing droplet assays in the presence of Cy5-labelled *in vitro* transcribed GFP RNA (800 nucleotides). As expected^45^, high RNA concentrations suppressed phase separation of WT FUS (Figures S1E and S1F). This inhibitory effect was slightly attenuated for RBdef FUS, consistent with its reduced RNA affinity. In contrast, CSdef FUS remained fully monodispersed across all tested RNA concentrations.

Together, these results establish that canonical RNA-binding and condensation are separable molecular activities of FUS that can be selectively impaired by point mutations.

### FUS condensation is required for its recruitment to paraspeckles and Cajal body formation

To investigate the cellular consequences of our FUS point mutations, we used the CRISPR-trap gene replacement approach^46^ to generate U2OS cell lines stably expressing eGFP-tagged WT, CSdef or RBdef FUS from the endogenous *FUS* locus (Figures 2A and 2B; Figures S2A and S2B). Previous reports have shown that mouse and human FUS are functionally interchangeable in DNA damage recruitment^17^, and fluorescent tagging (eGFP or mCherry) does not interfere with FUS activity in DNA repair^14^ or alternative splicing^41^. Using western blotting, we confirmed near endogenous expression levels of all eGFP-FUS constructs and verified the absence of endogenous FUS (Figure 2C). Immunofluorescence confirmed homogeneous expression and purity of the populations (Figure S2C). We observed a mild proliferation defect in the CSdef cells (Figures S2D and S2E), suggesting that condensation is required for FUS to promote cell growth^47^.

**Figure 2.**
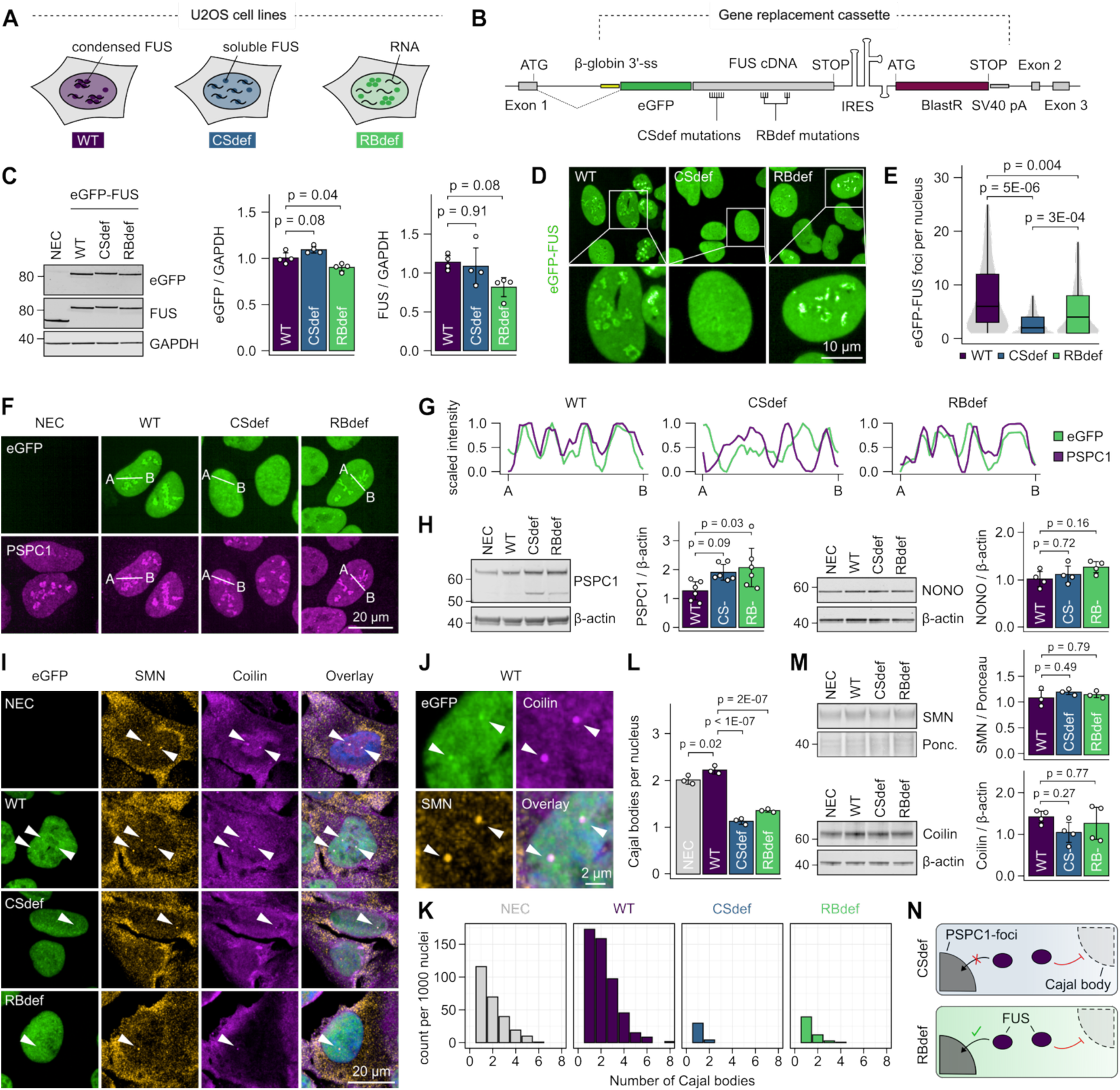
Condensation drives FUS recruitment and assembly of nuclear RNP condensates. (A) Schematic representation of the generated U2OS cell lines. (B) Genome-editing approach used to knock-in eGFP-FUS cDNA into the first intron of the endogenous *FUS* locus. (C) Western blot analysis of eGFP-FUS construct expression using anti-GFP and anti-FUS antibodies. GAPDH was used as a loading control. Bar plots show quantification of the eGFP signal relative to the mean across all conditions, and FUS signal relative to non-edited control (NEC). Data represent mean ± SD of N = 4 independent biological replicates. (D) Fluorescence microscopy images of eGFP-FUS in U2OS cell lines. Scale bar, 10 µm. (E) Violin-boxplot showing the number eGFP-FUS foci per foci-positive nucleus. The plot displays median lines, interquartile range (IQR) boxes and 1.5 × IQR whiskers. n = 1,485 (WT), 1,502 (CSdef), 1,681 (RBdef) nuclei, from N = 4 independent biological replicates. Statistical analysis was performed using the mean number of eGFP-FUS foci of each independent biological replicate, normalised to the mean across conditions. (F) Fluorescence microscopy images showing eGFP-FUS (green) and the paraspeckle marker PSPC1 (magenta). Scale bar, 20 µm. (G) Fluorescence intensity profiles of eGFP-FUS (green) and PSPC1 (magenta) along the white line indicated in the microscopy images. Intensities were normalised to the maximum value for each channel and scaled from 0 to 1. (H) Western blot analysis of PSPC1 (upper band) and NONO levels with β-actin as loading control. Bar plots show quantification relative to NEC. Data represent mean ± SD of N = 6 (PSPC1) or N = 4 (NONO) independent biological replicates. (I) Fluorescence microscopy images showing eGFP-FUS (green) and the Cajal body markers SMN (orange) and Coilin (magenta). Arrows indicate Cajal bodies. Scale bar, 20 µm. (J) Magnified view of two Cajal bodies in WT eGFP-FUS expressing cells from panel (I). (K) Bar plots showing the distribution of Cajal body number per 1000 Cajal body-positive nuclei. n = 309 (NEC), 596 (WT), 49 (CSdef), 82 (RBdef) nuclei, from N = 3 independent biological replicates. (L) Bar plots showing the mean number of Cajal bodies ± SD per positive nucleus from panel (K). (M) Western blot analysis of SMN and Coilin levels with β-actin or Ponceau S as loading control. Bar plots show quantification relative to NEC. Data represent mean ± SD of N = 3 (SMN) or N = 4 (Coilin) independent biological replicates. (N) Schematic model. FUS condensation promotes paraspeckle recruitment and Cajal body formation, whereas canonical RNA binding is required only for Cajal body formation. All statistical tests (C, E, H, L, M) were performed by one-way ANOVA followed by Tukey’s Honest Significant Difference (HSD) test for multiple comparison

As previously reported, WT eGFP-FUS localised predominantly to the nucleus and partitioned into micron-scale condensates (Figure 2D). RBdef FUS also formed condensates, albeit fewer, whereas CSdef FUS was diffusely nuclear. Automated nuclear segmentation and foci counting confirmed a marked reduction in the number of nuclear CSdef eGFP-FUS foci relative to WT (∼3-fold) alongside a modest reduction (∼1.5-fold) of RBdef eGFP-FUS foci (Figure 2E). To characterise the nature of these condensates, we examined co-localisation with known nuclear structures. We found that WT eGFP-FUS condensates co-localise with paraspeckle component 1 (PSPC1), a marker of paraspeckles which is also dynamically localises to perinucleolar caps upon transcription inhibition^48^ (Figure 2F). While RBdef eGFP-FUS was also enriched in PSPC1-positive foci, this localisation was lost in the CSdef mutant. This agrees with previous reports showing that the N-terminal low complexity domain^38^, and specifically tyrosine residues^39^, mediate FUS recruitment to paraspeckles. Quantification of PSPC1 foci revealed no change in CSdef, and a modest decrease in RBdef cells, mirroring the reduction in RBdef eGFP-FUS foci (Figure S2F). Notably, protein levels of PSPC1 were elevated (∼1.5-fold) in both mutants, and a truncated PSPC1 species appeared (Figure 2H), indicating perturbed paraspeckle homeostasis. Increased levels of PSPC1 were previously reported upon depletion of the core paraspeckle component non-POU domain-containing octamer-binding protein (NONO)^49^, which remained unchanged in our cell lines (Figure 2H). These findings suggest that FUS condensation is critical for its recruitment to PSPC1-containing paraspeckles and perinucleolar caps.

We next sought to evaluate the recruitment of our FUS mutants to Cajal bodies, given that FUS depletion or cytoplasmic mislocalisation impairs the formation of condensates containing Survival of Motor Neuron (SMN)^40,41^, a component of nuclear gems (without coilin) and Cajal bodies (with coilin). We found that WT eGFP-FUS was enriched, albeit mildly, in nuclear condensates positive for SMN and coilin, revealing their identity as Cajal bodies (Figures 2I and 2J). Strikingly, both the CSdef and RBdef mutants strongly impaired Cajal body formation (Figures 2K and 2L; Figure S2G). Protein levels of SMN and coilin remained unchanged (Figure 2M). We therefore conclude that, beyond FUS condensation, its ability to bind RNA provides an additional mechanistic layer that underlies Cajal body assembly or integrity (Figure 2N).

### FUS condensation and RNA-binding play distinct roles in the DNA damage response

FUS has been linked to multiple DNA repair pathways, including the repair of DNA double-strand breaks (DSBs), but the underlying mechanisms are only beginning to be understood. A well-characterised response of FUS is its rapid recruitment to sites of laser-induced DNA damage^13,14,17^. To evaluate the role of FUS condensation and canonical RNA-binding in this process, we subjected our U2OS cell lines to microirradiation with a 405 nm laser (Figure 3A). In line with previous reports, we observed fast recruitment of WT eGFP-FUS to DNA damage foci, reaching its peak approximately 50 seconds post-irradiation (Figures 3B and 3C). The focal nature of these compartments was particularly evident when microirradiation was focused on a single point instead of a larger region of interest (Figure S3A). Irradiated sites were enriched in histone H2AX phosphorylated at serine 139 (γH2AX), an early marker of DNA damage, and Poly(ADP-ribose) polymerase 1 (PARP1), which catalyses the addition of Poly(ADP-ribose) (PAR) chains to damaged chromatin (Figure S3B), confirming that the laser microirradiation generated *bona fide* DNA damage sites marked by early damage signalling and chromatin modification events. Recruitment of CSdef eGFP-FUS was significantly impaired (∼2-fold; Figures 3B and 3C), indicating that FUS condensation promotes its enrichment at DNA damage sites. This reconciles previous reports showing that the N-terminal low complexity domain promotes FUS enrichment after initial interactions between the FUS RGG domains and PAR chains^13,17^, which in turn promote FUS condensation *in vitro*^50^ in a tyrosine-dependent manner^51^. Further, it has been suggested that FUS also binds PAR through its RNA-binding interface within the RRM domain^52^. However, our RBdef mutant, which disrupts the relevant amino acids in this interface, was normally recruited to DNA damage sites (Figures 3B and 3C), arguing against a functional role of this interaction in FUS localisation.

**Figure 3.**
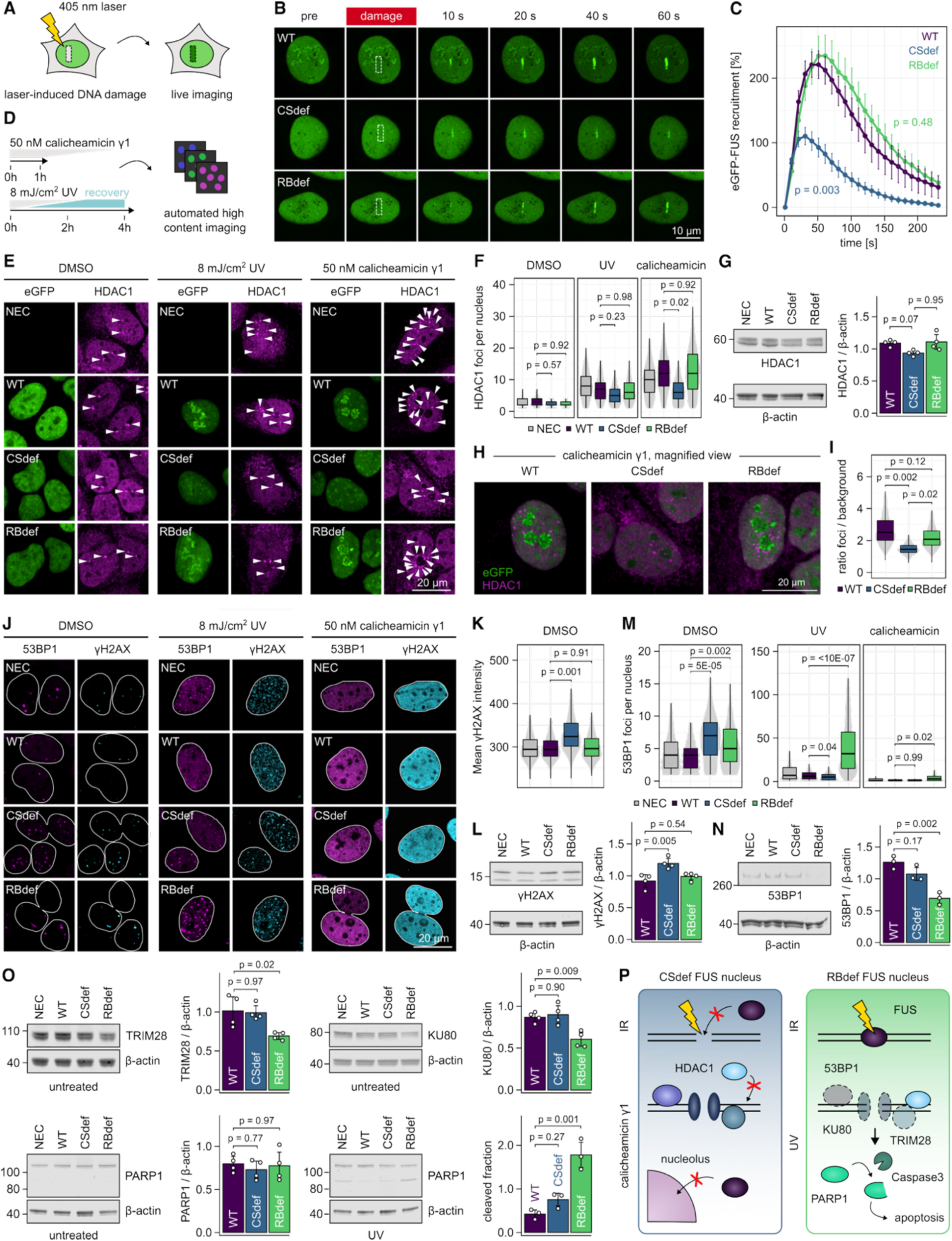
FUS condensation and RNA-binding distinctly contribute to FUS function in the DNA damage response. (A) Schematic of the laser microirradiation assay used to induce localised DNA damage. (B) Fluorescence microscopy images showing the localisation of eGFP-FUS before and at the indicated time points after DNA damage induction. Regions of interest (ROIs) targeted for microirradiation are outlined with dashed lines. Scale bar, 10 µm. (C) Quantification of eGFP-FUS recruitment to microirradiated sites, normalised to signal in a control ROI within the same nucleus. Data represent mean ± SEM of n = 12 (WT), 12 (CSdef), and 10 (RBdef) nuclei analysed from N = 3 independent biological replicates. For statistical analysis, recruitment was quantified as area under the curve, and differences between genotypes were assessed using a linear mixed-effects model with biological replicates as a random effect and pairwise comparisons via estimated marginal means. (D) Schematic of high-content imaging assays for DNA damage. (E) Fluorescence microscopy images showing the localisation of eGFP-FUS (green) and HDAC1 (magenta) in vehicle-treated cells (DMSO) and upon induction of DNA damage with UV or calicheamicin. White arrows indicate HDAC1 foci. Scale bar, 20 µm. (F) Violin-boxplots showing the number of HDAC1 foci per positive nucleus. DMSO-treated: n = 1275 (NEC), 1093 (WT), 1247 (CSdef), 1116 (RBdef); UV-treated: n = 719 (NEC), 819 (WT), 1,258 (CSdef), 826 (RBdef); calicheamicin-treated: n = 923 (NEC), 949 (WT), 1,426 (CSdef), 995 (RBdef) nuclei, from N = 3 independent biological replicates. (G) Western blot analysis of HDAC1 levels with β-actin as loading control. Bar plots show quantification relative to non-edited cells (NEC). Data represent mean ± SD of N = 3 independent biological replicates. (H) Magnified view of calicheamicin-treated cells from panel e, showing HDAC1 foci (magenta) and redistribution of eGFP-FUS (green) to perinucleolar structures. Scale bar, 20 µm. I) Violin-boxplot showing the enrichment of eGFP-FUS in the perinucleolar region relative to nucleoplasmic signal. n = 2,695 (WT), 4,537 (CSdef), 3,250 (RBdef) nuclei, from N = 3 independent biological replicates. (J) Fluorescence microscopy images of 53BP1 (magenta) and γH2AX (cyan) in vehicle-treated cells (DMSO) and upon induction of DNA damage with UV or calicheamicin. Nuclear outlines are defined by DAPI staining (white lines). Scale bar, 20 µm. (K) Violin-boxplot showing γH2AX intensity in DMSO-treated cells. n = 4,030 (NEC), 3,227 (WT), 3,464 (CSdef), 3,941 (RBdef) nuclei were analysed from N = 3 independent biological replicates. (L) Western blot analysis of γH2AX levels. Bar plots show quantification relative to NEC. Data represent mean ± SD of N = 4 independent biological replicates. (M) Violin-boxplot showing the number of 53BP1 foci per positive nucleus. DMSO: n = 3,566 (NEC), 2,738 (WT), 3,173 (CSdef), 3,657 (RBdef); UV: n = 1,054 (NEC), 1,248 (WT), 1,860 (CSdef), 1,770 (RBdef); calicheamicin: n = 451 (NEC), 447 (WT), 353 (CSdef), 1,099 (RBdef) nuclei from N = 3 independent biological replicates. (N, O) Western blot analysis of 53BP1, TRIM28, KU80 and PARP1 levels with β-actin as loading control. Bar plots show quantification relative to NEC. Data represent mean ± SD of N = 3 (53BP1, PARP1-UV) or N = 4 (TRIM28, KU80, PARP1-untreated) independent biological replicates. (P) Schematic model. Condensation enhances FUS recruitment to sites of laser-induced DNA damage, promotes HDAC1 foci formation and mediates re-distribution to perinucleolar structures upon severe DNA damage. In contrast, canonical RNA-binding by FUS maintains the expression of key NHEJ regulators, safeguarding effective DSB repair and preventing apoptotic progression. All violin-boxplots (F, I, K, M) display median lines, interquartile range (IQR) boxes and 1.5 × IQR whiskers. All statistical tests (F, G, I, K, M, L, N, O) were performed by one-way ANOVA followed by Tukey’s Honest Significant Difference (HSD) test for multiple comparisons.

Given the recruitment defect of our condensation-deficient mutant, we next asked whether FUS condensation also influences the recruitment of downstream DNA damage response (DDR) factors. Among the known damage-induced FUS interactors is the chromatin-modifying enzyme histone deacetylase 1 (HDAC1)^15^, which has been implicated in the cellular response to DSBs^42^. HDAC1 recruitment to DSB sites is reduced upon FUS depletion^15^. We therefore examined HDAC1 and eGFP-FUS localisation using automated high-content imaging following DNA damage induced either by the DNA strand-breaking agent calicheamicin or by exposure to UV light (Figure 3D). Both treatments led to an increase in HDAC1 foci formation, but this effect was impaired in CSdef cells (∼2-fold upon calicheamicin treatment; Figures 3E and 3F). HDAC1 protein levels were slightly reduced (93% of levels in unedited cells) in these cells, which may contribute to or exacerbate the impaired foci formation (Figure 3G). While we did not detect an enrichment of eGFP-FUS in HDAC1 condensates, we observed a striking redistribution of eGFP-FUS to nucleolar structures following calicheamicin or UV treatment (Figure 3H). This nucleolar translocation was significantly impaired (∼2-fold) in CSdef cells (Figure 3I). Notably, this translocation defect is not linked to altered FUS phosphorylation, as all eGFP-FUS constructs exhibited the characteristic shift to higher apparent molecular weight upon calicheamicin treatment (Figure S3C)^53^. Nucleolar localisation of FUS has been previously reported in response to DNA damage and is associated with repression of RNA polymerase II transcription^54^. We did not observe this nucleolar localisation of FUS following treatment with other DNA damage-inducing agents, such as the topoisomerase inhibitors camptothecin or etoposide, which induced lower amounts of DNA damage (Figures S3D and S3E). This suggests that translocation of FUS to the nucleolus is a feature associated with severe genotoxic stress.

To further characterise the DSB response in our system, we examined two canonical DDR markers: The early damage sensor γH2AX and p53-binding protein 1 (53BP1), a downstream mediator that promotes non-homologous end joining (NHEJ). Previous studies reported increased γH2AX levels following nuclear FUS depletion^14,55^. In agreement, we observed a modest increase (∼1.2-fold) in γH2AX signal under basal conditions in CSdef eGFP-FUS expressing cells, while RBdef eGFP-FUS expressing cells showed no change (Figures 3J and 3K). These findings were confirmed by western blot analysis (Figure 3L). As expected, treatment with calicheamicin or UV light induced a robust increase in nuclear γH2AX levels and 53BP1 foci formation (Figure 3J), while camptothecin and etoposide elicited a similar, slightly milder, response (Figures S3E-H). eGFP-FUS was not enriched in γH2AX or 53BP1 foci (Figure S3F). Unexpectedly, UV-treated RBdef cells exhibited a striking increase in 53BP1 foci relative to WT (∼5-fold) or unedited cells (Figure 3M). This phenotype was also observed upon treatment with camptothecin or etoposide (Figure S3I), suggesting that RNA-binding by FUS influences the efficiency or resolution of DSB repair via the NHEJ pathway.

To investigate the molecular basis for this defect, we quantified protein levels of 53BP1 as well as several NHEJ-associated factors, informed by transcriptomic data (see below). In RBdef cells, we observed significantly decreased levels of both 53BP1 (∼1.8 fold) and X-ray repair cross-complementing protein 5 (∼1.4 fold; XRCC5, commonly referred to as Ku80), a DNA end-binding protein critical for early double-strand break recognition (Figures 3N and 3O). These downregulations persisted upon UV exposure (Figure S3J). In addition, levels of tripartite motif-containing 28 (TRIM28, also known as KAP1), a chromatin scaffold protein that promotes chromatin relaxation around DSB sites, were also decreased under basal (∼1.5-fold) and damage-induced (∼1.7-fold) conditions (Figure 3O; Figure S3J). These molecular deficits were accompanied by a marked increase in the cleavage of poly(ADP-ribose) polymerase 1 (PARP1) in UV-treated RBdef cells (∼4.2-fold), a hallmark of apoptotic signalling induced by unresolved DNA damage^56,57^.

In summary, our findings indicate that FUS condensation facilitates robust recruitment to DNA damage sites, supports HDAC1 foci formation, and mediates nucleolar translocation upon severe genotoxic stress. In contrast, canonical RNA-binding by FUS is essential for maintaining the expression of key NHEJ regulators, including 53BP1, Ku80 and TRIM28, thereby safeguarding effective DSB repair and preventing apoptotic progression (Figure 3P).

### The RNA-binding landscape of FUS is shaped by condensation and canonical RNA-binding

How canonical RNA-binding and condensation of FUS affect its transcriptome-wide RNA-binding landscape remains unknown. To address this, we subjected our eGFP-FUS constructs to a streamlined version of individual nucleotide-resolution crosslinking and immunoprecipitation (siCLIP) alongside total RNA sequencing of our engineered U2OS lines. Upon purification with GFP nanobodies, we detected strictly crosslink- and RNase-dependent protein-RNA complexes migrating slower than free eGFP-FUS (Figure S4A). The final siCLIP experiments were performed in triplicates (Figure 4A). As expected, the signal of the protein-RNA complexes was reduced for RBdef eGFP-FUS (∼2-fold; Figure 4B), with residual signal pointing towards transient, low-affinity binding events or interactions mediated through the RGG domains. RNA was isolated from the regions indicated by the dashed lines and converted into cDNA libraries for high-throughput sequencing.

**Figure 4.**
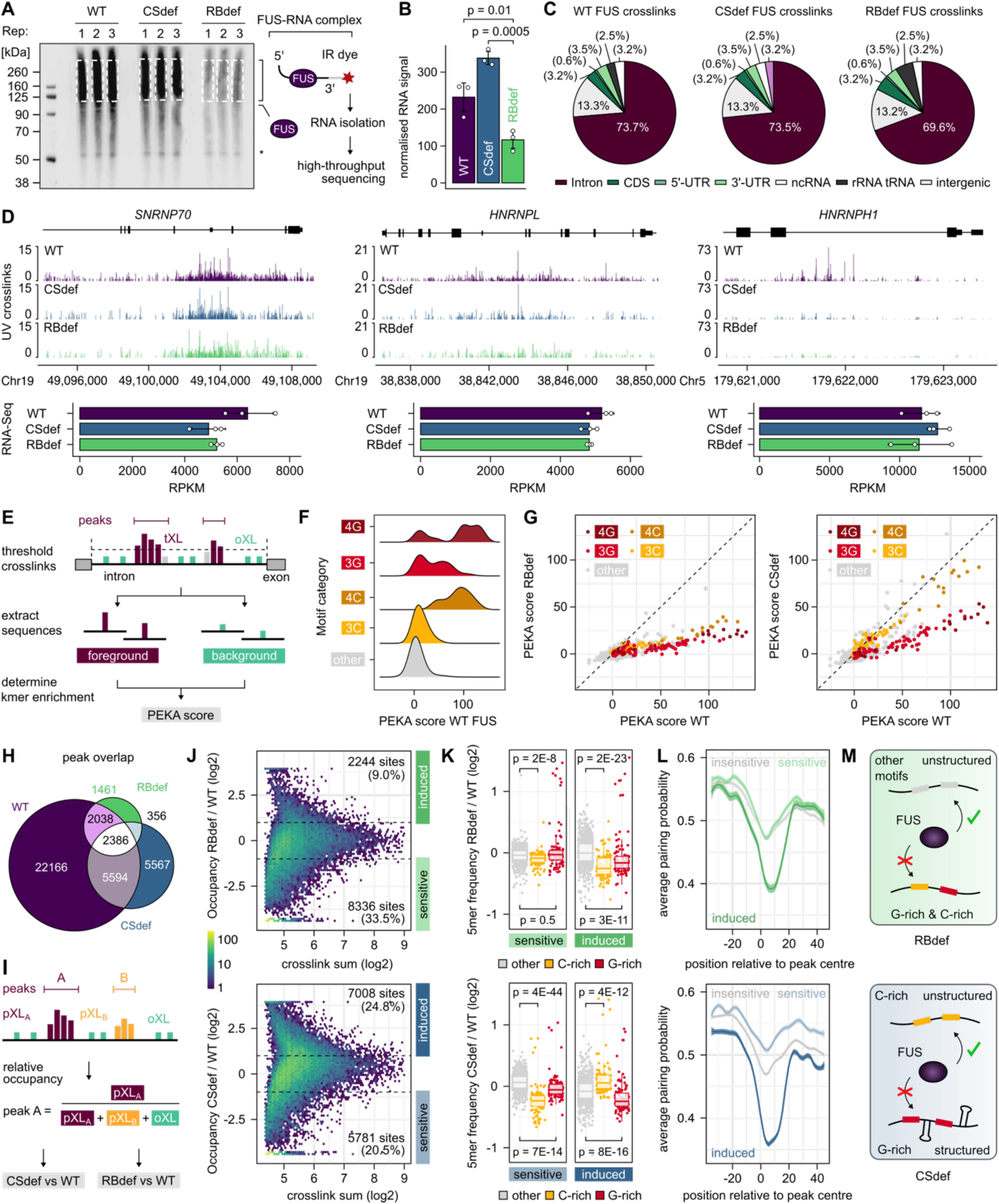
The RNA-binding landscape of FUS is shaped by condensation and canonical RNA-binding. (A) Visualisation of protein-RNA complexes formed by eGFP-FUS following SDS-PAGE and transfer onto a nitrocellulose membrane. Crosslinked RNA fragments are 3’-labelled with a Cy5-labelled adapter. Dashed boxes indicate regions that were excised for RNA isolation. (B) Bar plot showing the RNA signal in the dashed boxes normalised to the amount of purified eGFP-FUS protein. Data represent mean ± SD of N = 3 independent biological replicates. Statistical significance was analysed by one-way ANOVA followed by Tukey’s HSD test for multiple comparisons. (C) Pie chart summarising the fraction of crosslink events within different genomic regions. n = 18,284,974 (WT), 7,776,767 (CSdef), 6,203,397 (RBdef). (D) Genome browser view of the *SNRNP70*, *HNRNPL* and *HNHNPH1* genes showing the positions of UV crosslinks detected by eGFP-FUS siCLIP. Bar plots show the expression levels of these genes determined by RNA-Seq as reads per kilobase of transcript per million mapped reads (RPKM). Data represent mean ± SD of N = 3 independent biological replicates. (E) Schematic representation of the positionally enriched k-mer analysis (PEKA). Sequences around thresholded crosslink (tXL) sites within peaks are used as foreground and compared to background sequences from reference crosslink sites located outside of peaks (oXLs), accounting for intrinsic UV crosslink biases. (F) Ridgeline plot showing the distribution of PEKA scores for 5-mers containing ≥ 4 guanosines (4G, dark red), 3 guanosines (3G, red), ≥ 4 cytosines (4C, dark orange), 3 cytosines (3C, orange) or the remaining 5-mers (other, grey) for WT eGFP-FUS. (G) Scatterplots showing the correlation of PEKA scores for each 5-mer between WT and RBdef as well as WT and CSdef eGFP-FUS. G-rich and C-rich 5-mers are coloured as in panel (F). (H) Venn diagram showing the overlap of high-confidence peaks between the different eGFP-FUS constructs. (I) Schematic representation of the binding site occupancy analysis. For each high-confidence peak, the sum of crosslink sites was normalised to the total number of crosslinks in the pre-mRNA to account for differences in library size and transcript levels. (J) Hexagonal binning plot showing the ratio of eGFP-FUS occupancy against the sum of crosslink events for each binding site, based on the combined set of high-confidence peaks from both conditions. The colour scale indicates point density. Dashed lines mark two-fold changes in either direction. (K) Boxplots showing the ratios of G-rich (red), C-rich (orange), and other (grey) 5-mer frequencies in binding sites affected by the condensation or RNA-binding mutations, with individual 5-mers represented as jittered points. Differences in the distribution of 5-mer frequency ratios between motif classes were assessed using a two-sided Wilcoxon rank-sum (Mann-Whitney U) test. (L) Base-pairing probabilities were computed using the 100 nucleotide RNA regions surrounding binding sites affected by the condensation or RNA-binding mutations. Line plots display base-pairing probability profiles from positions -35 to +45 relative to peak centre. (M) Schematic model. Condensation promotes FUS binding to G-rich and structured binding sites but is dispensable for binding to C-rich motifs. In contrast, canonical RNA-binding confers specificity towards both G-rich and C-rich motifs.

In agreement with previous iCLIP studies, WT eGFP-FUS exhibited widespread binding across nascent transcripts, with ∼75% of crosslink events mapping to introns (Figure 4C). This regional specificity was preserved in the CSdef and RBdef mutants. However, while binding of FUS to some introns, such as a conserved intron in the SNRNP70 pre-mRNA, was relatively unaffected by the mutations, other introns (e.g. in the HNRNPL and HNRNPH1 pre-mRNAs) displayed clear signs of RNA-binding- and/or condensation-dependency (Figure 4D).

We therefore evaluated how the point mutations affect the sequence-specificity of FUS. Hereto, we quantified the prevalence of pentanucleotide (5-mer) motifs around intronic crosslink clusters using positionally enriched k-mer analysis (PEKA) (Figure 4E). Among the most significantly enriched 5-mers for WT eGFP-FUS, we found a strong overrepresentation of G-rich motifs containing GGA or GGU (Figures 4F; Figures S4B and S4C), in line with the reported sequence-specificity of the FUS zinc finger domain. Surprisingly, C-rich motifs were also strongly overrepresented among the top 5-mers, hinting at a previously unrecognised sequence-specificity of FUS. While C-rich motifs were enriched very close to the crosslink sites, G-rich motifs exhibited a broader positional distribution, suggestive of multivalent binding (Figure S4D). To determine whether this enrichment of G- and C-rich motifs depends on functional RNA-binding domains, we compared the PEKA scores of WT and RBdef eGFP-FUS. Indeed, we found that these scores were globally reduced for the RBdef mutant, confirming that the RRM and ZnF domains confer sequence-specificity to G-rich and C-rich motifs (Figure 4G). For the CSdef mutant, we observed an unexpected motif-specific effect: While C-rich motif enrichment was largely preserved, G-rich motifs showed a pronounced reduction in PEKA scores relative to WT.

In line with altered sequence-specificity, high-confidence crosslink clusters (hereafter referred to as ‘binding sites’) showed only modest overlap between the different FUS constructs (Figure 4H). This prompted us to further investigate how the CSdef and RBdef mutations affect binding at the level of whole transcripts and individual binding sites. Hereto, we quantified and normalised the occupancy of both mutant proteins (adjusted for library size and transcript abundance) on protein-coding transcripts and individual FUS binding sites (Figure 4I). While both mutants showed only modest differential binding at the transcript level (Figure S4E), more pronounced differential binding emerged at the level of individual binding sites: For RBdef eGFP-FUS, ∼34% of binding sites showed > 2-fold reduced occupancy, whereas only ∼9% of binding sites showed increased occupancy (Figure 4J). Sites with increased RBdef eGFP-FUS binding were depleted in G-rich and C-rich 5-mers (Figure 4K; Figure S4F) and exhibited lower predicted RNA secondary structure propensity (Figure 4L), suggesting that loss of canonical RNA-binding promotes binding to atypical, non-cognate sites in unstructured regions. In contrast, CSdef eGFP-FUS displayed increased occupancy at ∼25% and decreased occupancy at ∼21% of binding sites (Figure 4J). Whereas CSdef-sensitive sites (decreased binding) were depleted of C-rich motifs and showed higher secondary structure propensity, CSdef-induced sites (increased binding) were enriched in C-rich motifs (Figure 4K; Figure S4G) and displayed reduced secondary structure propensity (Figure 4L). This corroborates the finding that condensation-deficient FUS retains its preference for C-rich motifs.

In summary, our data indicate that the canonical RNA-binding domains of FUS confer specificity to G-rich and C-rich motifs. While FUS condensation is dispensable for efficient binding to C-rich motifs, it promotes binding to G-rich motifs and structured RNA elements (Figure 4M).

### RNA-binding and condensation by FUS regulate transcription of ion channel mRNAs

We next sought to identify downstream consequences of altered RNA-binding by CSdef and RBdef FUS on FUS-dependent gene regulation. Principal component analysis confirmed clustering of the biological replicates (Figure S5A). Differential expression analysis using DESeq2 identified 1,326 downregulated genes and 889 upregulated genes (fold change > 1.5 and adjusted p-value < 1E-05) in RBdef eGFP-FUS cells and 1,534 downregulated and 1,221 upregulated genes in CSdef eGFP-FUS cells (Figure 5A; Table S1). These changes primarily affected protein-coding genes (Figure S5B) and showed only modest overlap between the mutant cell lines (Figures 5B and S5C). Nevertheless, global expression changes were significantly correlated (ρ = 0.44; Figure 5C), suggesting that FUS condensation and RNA-binding often regulate gene expression in a cooperative manner.

**Figure 5.**
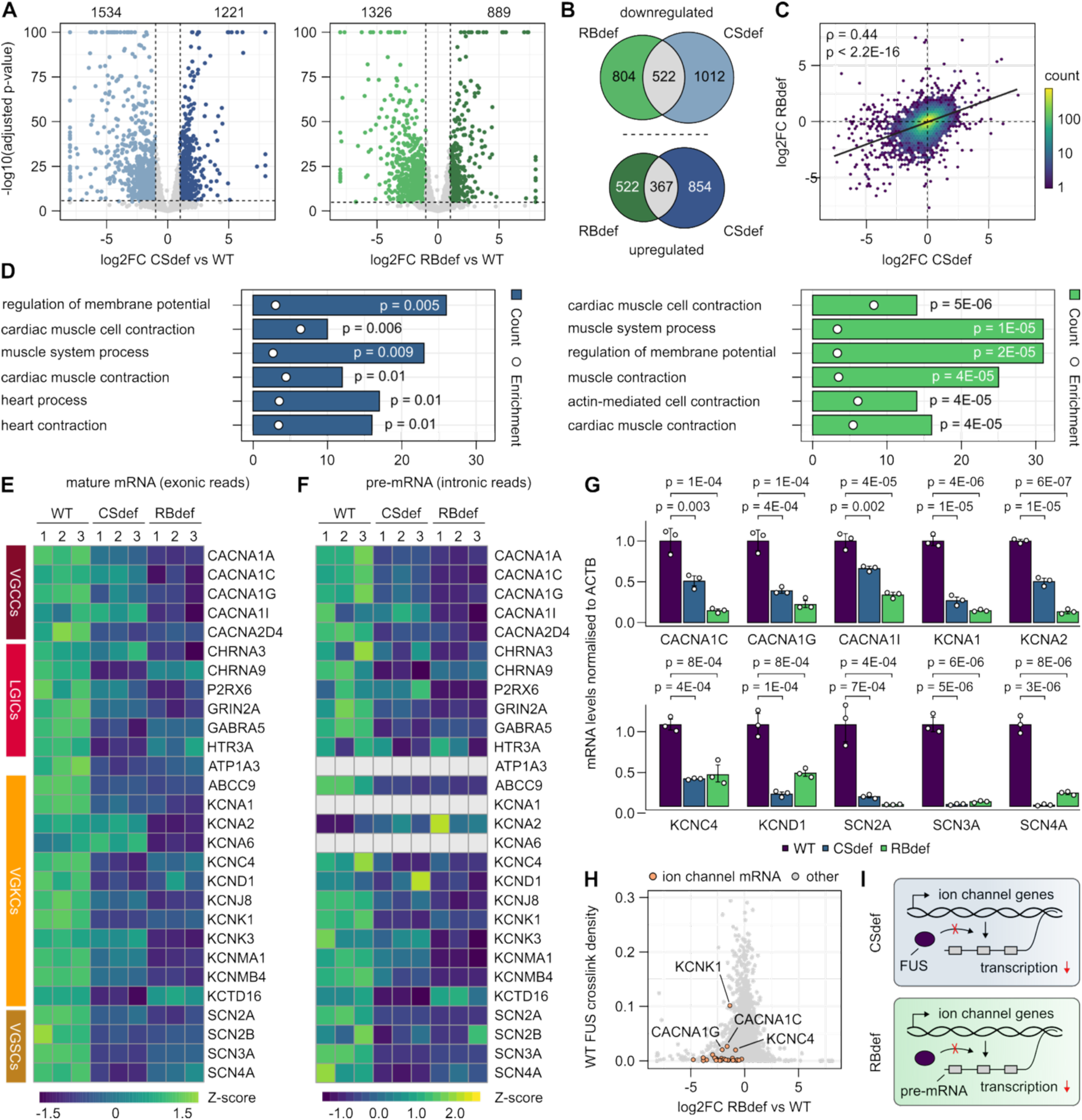
FUS condensation and RNA-binding contribute to ion channel mRNA expression. (A) Volcano plots showing differential transcript abundance between mutant and WT cell lines analysed by DESeq2 from N = 3 independent biological replicates. Vertical dashed lines indicate 1.5-fold change, horizontal lines mark the significance threshold (adjusted p-value < 10E-05). Counts of differentially up-and downregulated genes are indicated. (B) Venn diagrams showing the overlap of up- and downregulated genes between CSdef and RBdef cells relative to WT. (C) Hexagonal binning plot showing the correlation of transcript levels between CSdef and RBdef cells relative to WT. The black line shows a linear regression fit. Spearman correlation (ρ) and p-value are indicated. (D) Gene Ontology (GO) enrichment analysis of the 500 most downregulated genes in CSdef and RBdef cells relative to WT. Analysis was performed using the Biological Process (BP) ontology in clusterProfiler with Benjamini–Hochberg correction for multiple testing (adjusted p-value < 0.05). Significantly enriched GO terms are shown, ranked by adjusted p-value. (E) Heatmap showing expression levels of ion channel mRNAs (based on exonic reads) across cell lines, grouped by category: voltage-gated calcium channels (VGCCs), ligand-gated ion channels (LGICs), voltage-gated potassium channels (VGKS), and voltage-gated sodium channels (VCSCs). Expression values represent SD from mean of variance stabilised values across rows. Upregulated genes are shown in green, downregulated genes are shown in purple. (F) Heatmap showing expression levels of ion channel pre-mRNAs (based on intronic reads) across cell lines. Expression values represent SD from mean of reads per kilobase of transcript, per million mapped reads (RPKM) values across rows. (G) RT-qPCR validation of selected ion channel mRNAs. Data represent mean relative abundance normalised to the WT mean, with error bars showing standard deviation from N = 3 independent biological replicates. Statistical significance was computed using ΔΔCt values by one-way ANOVA followed by Tukey’s Honest Significant Difference (HSD) test for multiple comparisons. (H) Scatterplot showing the correlation between WT eGFP-FUS crosslink density and changes in transcript abundance in RBdef relative to WT cells. Ion channel mRNAs from panel (F) are highlighted in orange and the four ion channel mRNAs with the highest crosslink densities are labelled. (I) Schematic model. FUS condensation and RNA-binding contribute to the transcription of ion channel pre-mRNAs independently of direct FUS binding to these transcripts.

Using GO term enrichment analysis, we found that genes linked to the regulation of membrane potential and muscle contraction were significantly overrepresented among the 500 most downregulated genes in both CSdef and RBdef eGFP-FUS cells (Figure 5D). This enrichment was primarily driven by the downregulation of mRNAs encoding ion channel subunits, particularly members of the voltage-gated calcium channels (*CACNA* genes), potassium channels (*KCNA* genes) and sodium channels (*SCN* genes) (Figure 5E). The downregulation was also evident at the pre-mRNA level, indicating that reduced transcription, rather than aberrant RNA processing, underlies the observed downregulation (Figure 5F; Figure S5D). Using RT-qPCR, we independently validated the downregulation of 10 selected ion channel mRNAs (Figure 5G), achieving a 100% confirmation rate and thereby supporting the robustness of the RNA-Seq data. Finally, we questioned whether direct FUS - RNA interactions could promote transcription of ion channel mRNAs. However, the eGFP-FUS crosslink density on these transcripts was low (Figure 5H), arguing against a significant role of direct RNA-binding to the effect.

In summary, our findings suggest that condensation and canonical RNA-binding by FUS influence the transcription of ion-channel mRNAs independently of extensive direct binding to these transcripts (Figure 5I).

### RNA-binding and condensation cooperatively modulate FUS-dependent alternative splicing

We finally aimed to dissect the roles of RNA-binding and condensation in FUS-dependent alternative splicing. Using rMATS, we identified 620 affected splicing events in CSdef cells and 407 splicing events in RBdef cells (Figures 6A and 6B; Figure S6A; Table S2). In both mutants, skipped exons were the most prevalent type of splicing alteration, with a consistent bias toward reduced exon inclusion compared to WT cells. Conversely, intron retention, the second most common type of splicing alteration, showed a consistent bias toward increased intron retention in mutant cells. As with the observed changes in gene expression, these alternative splicing changes were modestly but significantly correlated (Figure 6C), suggesting that FUS condensation and RNA-binding often function cooperatively to regulate RNA processing.

**Figure 6.**
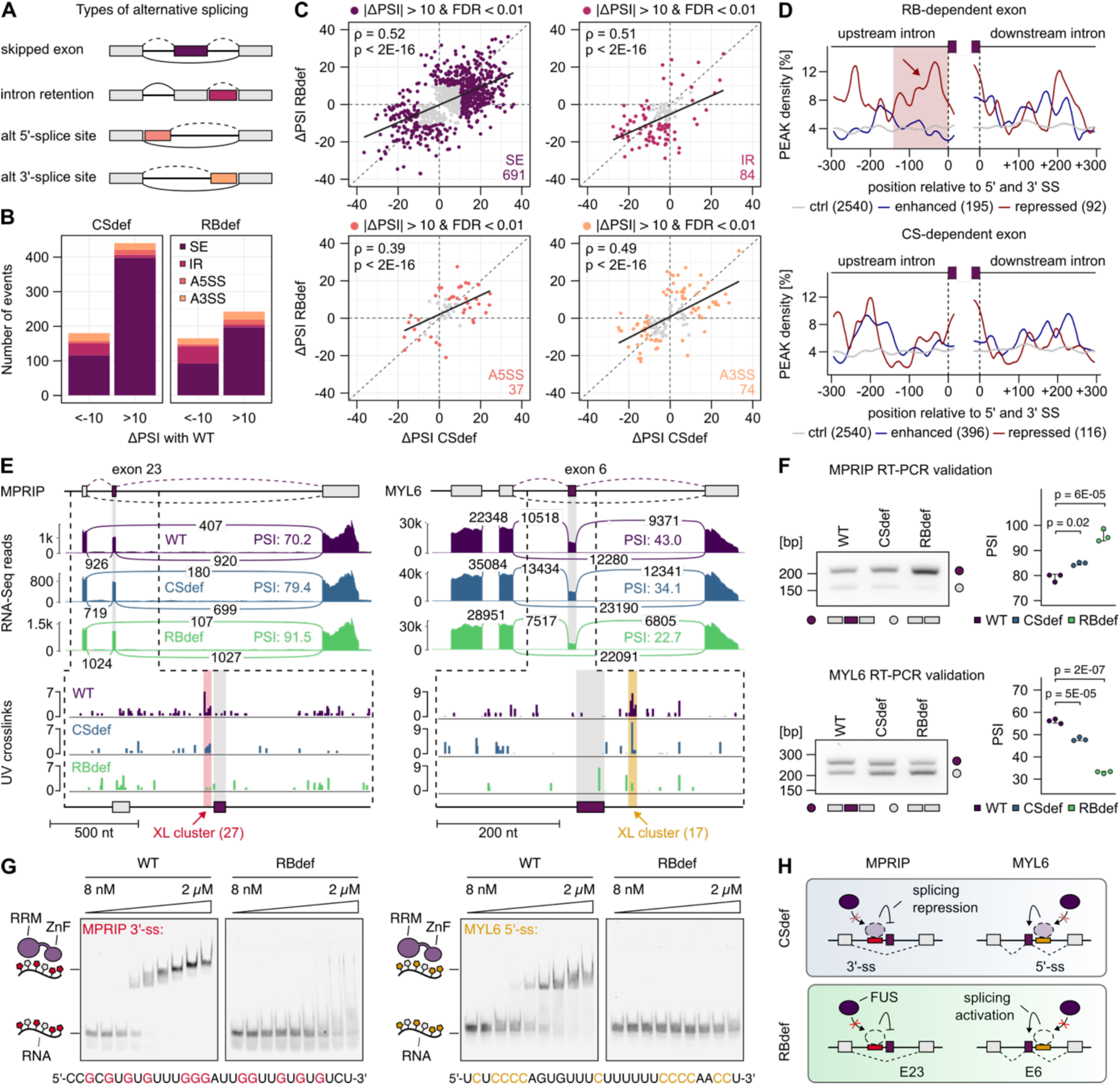
FUS condensation and RNA-binding co-operatively regulate alternative splicing. (A) Schematic representation of the types of alternative splicing analysed in this study. (B) Stacked bar plots summarising the number of significant splicing alterations detected by rMATS, defined as absolute change in percent spliced in (|ΔPSI|) > 10 and false discovery rate (FDR) < 0.01. Colours indicate the type of alternative splicing event: skipped exon (SE), intron retention (IR), alternative 5’-splice site (A5SS), and alternative 3’-splice site (A3SS). (C) Scatterplots showing the correlation of ΔPSI values from significant alterations between CSdef and RBdef cells relative to WT, grouped by type of alternative splicing event. The black line shows a linear regression fit. Spearman correlation (ρ) and p-value are indicated. (D) RNA maps showing normalised density of WT eGFP-FUS crosslink peaks relative to the 5’- and 3’-splice sites of repressed (red), promoted (blue) and control (grey) cassette exons identified in CSdef and RBdef cells. (E) Genome browser view of *MPRIP* and *MYL6* regions with FUS-regulated alternative exons. Top: Sashimi plots showing RNA-Seq junction reads and PSI values. Bottom: Bar plots of eGFP-FUS crosslinks around regulated exons, with clusters highlighted in colour. (F) RT-PCR validation of alternative splicing in *MPRIP* and *MYL6*. Dot plots show PSI values (Mean ± SD of N = 3 independent biological replicates. Statistical significance was analysed by one-way ANOVA followed by Tukey’s HSD test for multiple comparisons. (G) Electrophoretic mobility shift assays (EMSAs) with GB1-tagged WT or RBdef FUS (amino acids 269-454) and Cy5-labelled RNA derived from the crosslink clusters near regulated exons in *MPRIP* and *MYL6*. (H) Schematic model. Canonical RNA-binding and, to a lesser extent, condensation promote binding of FUS to G-rich (red) and C-rich (orange) binding sites near alternative exons in *MPRIP* and *MYL6* to regulate exon inclusion.

To determine whether altered exon inclusion can be explained by direct FUS binding, we generated RNA maps that visualise the density of WT eGFP-FUS crosslink clusters in or around RNA-binding and condensation-dependent exons. In this meta-analysis, we observed a clear enrichment of FUS binding immediately upstream of exons that are repressed in an RNA-binding dependent manner (Figure 5D), indicating that binding of FUS at the 3’-splice site typically prevents exon recognition. In contrast, we only detected a mild enrichment of FUS binding near condensation-sensitive exons and no clear positional logic. This suggests that a subset of events is regulated indirectly, for example by FUS regulating other RNA-binding proteins or maintaining Cajal bodies.

To further investigate splicing alterations that can be explained by direct FUS binding, we therefore turned our attention to individual pre-mRNAs. Specifically, we examined two pre-mRNAs, MPRIP and MYL6, where changes in exon inclusion coincide and correlate with altered binding of the CSdef and RBdef mutants to G-rich (MPRIP) or C-rich (MYL6) motifs near the regulated exon (Figure 6E). Similar patterns were also observed for other transcripts where splicing differences are accompanied by reduced binding to G-rich (EIF4H, PCBP2) or C-rich binding sites (TANGO2) (Figure S6B). Using RT-PCR, we independently validated these splicing alterations, confirming the robustness of the data (Figure 6F; Figure S6C).

To validate FUS binding to specific sites near regulated exons, we performed electrophoretic mobility shift assays using a GB1-tagged minimal FUS-RBD construct encompassing the RRM, RGG2 and the ZnF domains (aa 269 - 447) and Cy5-labelled RNA oligonucleotides derived from the G-rich motif cluster upstream of MPRIP exon 23 and the C-rich motif cluster downstream of MYL6 exon 6. Both RNAs were bound by FUS-RBD at protein concentrations above 62 nM (MPRIP) and 125 nM (MYL6) (Figure 6G). This binding was almost completely abolished by the RBdef mutations, confirming that canonical RNA-binding confers specificity to these motifs.

In summary, our data indicate that condensation and canonical RNA-binding cooperatively regulate alternative splicing, particularly exon inclusion. This is exemplified by individual events (e.g. MPRIP and MYL6) where different splicing outcomes are directly associated with altered binding to G-rich or C-rich binding sites near regulated exons, indicating that FUS leverages both sequence preferences to regulate alternative splicing (Figure 6H).

## Discussion

In this study, we set out to dissect how condensation and canonical RNA-binding contribute to the physiological functions of FUS. Using only a small number of precisely targeted point mutations, avoiding the confounding effects often associated with more invasive perturbations like deletions, we uncoupled these two biochemical properties and thereby uncovered previously hidden layers of regulation that cannot be resolved with traditional loss-of-function studies. Together, these findings provide a mechanistic framework for understanding how FUS integrates its condensation capacity with sequence-specific RNA recognition to coordinate nuclear organisation, RNA metabolism, and genome stability. Notably, this separation-of-function approach offers a conceptual paradigm applicable to the wide array of condensate-forming RBPs and provides a general strategy for disentangling the intertwined contributions of phase behaviour and RNA recognition to their molecular functions. With growing interest in targeting RBP condensation in neurodegenerative disease, delineating these contributions will be critical to anticipate the outcomes of such interventions, and mitigate potential adverse effects.

### Dual layers of FUS regulation in nuclear condensate organisation and the DNA damage response

Our findings demonstrate that FUS condensation plays a key role in enabling FUS to partition into membraneless compartments such as paraspeckles, perinucleolar caps, Cajal bodies and DNA damage foci. This aligns with the conceptualisation of these structures as biomolecular condensates formed through phase separation of their constituents^35^, and suggests that FUS recruitment to each structure is governed by shared biophysical principles. While we cannot formally exclude the possibility that the small number of mutated residues within the low-complexity domain might also influence recruitment via protein-protein interactions, a FUS mutant in which 12 prolines were substituted with alanines in the same region, selectively impairing fibril formation rather than condensation, displayed normal localisation to paraspeckles, perinucleolar caps and DNA damage foci^58^. This observation suggests that the observed changes for CSdef FUS are most likely a direct consequence of altered condensation capacity. While canonical RNA-binding is largely dispensable for FUS recruitment to these condensates, we discovered that is strictly required for Cajal body formation, revealing an unanticipated mechanistic layer that underlies the formation of these structures. We propose two, non-mutually exclusive, models: canonical RNA-binding may stabilise the dynamic molecular network that maintains Cajal bodies, or it may act upstream, by recruiting specific RNA substrates or RNP complexes that nucleate Cajal body assembly. This idea is supported by our previous finding that FUS directly binds small nuclear RNAs^43^, which transit through Cajal bodies for maturation and recycling^37^.

In line with results from deletion mutants^13^, we show that FUS condensation enhances its recruitment to DNA damage sites, leading to increased basal levels of γH2AX and a reduction of HDAC1 recruitment to DNA damage foci in condensation-deficient cells. These results provide the first direct evidence that FUS condensation actively promotes these processes. However, a second mechanistic layer also emerged for the role of FUS in the DNA damage response, where the most pronounced defect in DNA repair (increased number of 53BP1 foci and apoptotic progression) is linked to the canonical RNA-binding activity of FUS, likely mediated through its role in maintaining the expression of key NHEJ regulators, including 53BP1, Ku80 and TRIM28. This highlights the value of precise separation-of-function mutations, which make it possible to distinguish direct molecular roles from broader, indirect regulatory effects and thereby advance our understanding of how multifunctional proteins like FUS safeguard genome stability.

### The canonical RNA-binding domains of FUS confer specificity to G-rich and C-rich motifs

Previous crosslinking and immunoprecipitation studies have concluded that FUS binds nascent pre-mRNAs in a widespread manner, with only a mild preference for G-rich motifs like GGU or GUGGU^10,11^. Our study establishes that the canonical RNA-binding domains are not required for the bulk of this widespread RNA-binding, supporting the view that the RGG domains are major contributors to FUS RNA-binding^24–26^, likely through direct, non-specific electrostatic interactions. Instead, the RRM and ZnF domains confer specificity to a subset of binding sites enriched in G-rich motifs and, unexpectedly, C-rich motifs. While the ZnF is known to bind GGU motifs through extensive hydrogen bonding and aromatic stacking interactions with the two guanosines^23^, it remains unclear which domain mediates the previously unrecognised preference for C-rich motifs. The RRM can accommodate RNA stem loops but also exhibits relatively broad specificity for single-stranded sequences. Intriguingly, the RRM binds to ACGCGC with 4- to 5-fold higher affinity than AAUAAA^23^, suggesting that the presence of cytosines enhances binding specificity and/or affinity, though this hypothesis requires experimental validation.

### FUS condensation promotes binding to G-rich, structured RNA sequences

RNA affects the condensation behaviour of RBPs^45,59–63^, yet our understanding of how condensation fine-tunes the transcriptome-wide RNA-binding landscape of RBPs remains remarkably limited. A seminal study showed that condensation of the ALS-linked protein TDP-43 is required for efficient binding to long (>100 nt) RNA regions with dispersed, low-affinity motifs but is dispensable for binding to shorter regions dense in high-affinity motifs^64^. In this context, condensation amplifies individually weak interactions across extended regions to stabilise RNA engagement. By contrast, our results show that the condensation capacity of FUS is required for efficient binding to G-rich, structured RNA sequences, but not for interaction with C-rich motifs and linear RNA sequences. We therefore propose that FUS employs condensation to overcome structural, rather than affinity-based obstacles. G-rich motifs can form stable secondary structures, including G-quadruplexes, which may present binding sites that are sterically occluded and hence require condensation to increase local FUS concentration and/or enable multivalent engagement by multiple FUS molecules. By contrast, C-rich motifs and unstructured RNA sequences present more accessible, linear binding surfaces that can be engaged effectively by individual FUS molecules without the need for higher-order assembly. Interestingly, G-rich RNAs with propensity to form G-quadruplexes are themselves intrinsically more prone to phase separation than C-rich RNAs^65–67^. This inherent self-association may synergise with FUS condensation, such that structured G-rich RNA sequences act not only as condensation-dependent binding targets but also as co-nucleators of higher-order assemblies. Future *in vitro* reconstitution experiments using defined RNA substrates with tuneable secondary structure and controlled phase-separation conditions could help dissect the mechanisms by which FUS condensation enhances binding to G-rich motifs within structured contexts.

Together, these studies suggest a unifying principle: condensation acts as a context-dependent amplifier of RNA-binding to specific sites within an RNA molecule, instead of promoting binding to whole transcripts. In doing so, condensation enables RBPs to overcome distinct barriers - sparse motif clustering for TDP-43 and structural inaccessibility for FUS - thereby expanding the repertoire of RNA substrates they can recognise and regulate.

### Synergistic roles of FUS condensation and RNA-binding in regulating gene expression

Depletion of FUS causes widespread alterations in gene expression, impacting hundreds of genes through both transcriptional regulation and RNA processing^8–11,68^. Our study shows that FUS condensation plays a key role in both functions and often acts synergistically with canonical RNA-binding.

At the transcriptional level, both condensation and RNA-binding are required to sustain ion channel gene expression, a function that appears largely independent of direct FUS interaction with the corresponding pre-mRNAs. This points to an upstream role for FUS in scaffolding regulatory condensates at chromatin. Supporting this view, FUS occupies hundreds of promoters^69^, engages transcription factors^70–72^ and RNA polymerase II^73^, and forms transcription-associated condensates^27,74^. The canonical RNA-binding domains may stabilise these assemblies via promoter-associated non-coding RNAs^8,75^ or DNA^76^. Our point mutants provide powerful tools to dissect the contributions of condensation and RNA-binding to chromatin engagement and transcriptional regulation, for example through ChIP-based approaches.

By contrast, regulation of alternative splicing displayed a more direct relationship to RNA-binding. Here, FUS binding is modestly enriched in the vicinity of regulated exons, most notably at the 3’-splice site of exons repressed in an RNA-binding dependent manner. This enrichment indicates that FUS can act directly on splicing regulation, with a tendency toward repression, consistent with previous findings^10^. Building on this global trend, our use of targeted FUS point mutants enabled us to pinpoint specific splicing events explained by altered FUS binding to both G- and C-rich motifs, indicating that FUS exploits both sequence preferences for regulatory purpose. These include events (e.g. EIF4H and MPRIP) that have also been detected in published datasets acquired upon FUS depletion in human as well as mouse systems^68^, indicative of conserved regulation.

In conclusion, our findings establish FUS as a multifunctional regulator whose condensation and sequence-specific RNA-binding act in concert to control nuclear condensate organisation, RNA metabolism, and genome stability, providing a framework for how RBPs integrate biophysical and biochemical properties to shape their physiological functions. Looking ahead, our rigorously characterised mutants will serve as powerful tools to dissect the contribution of FUS condensation and RNA-binding to ALS pathogenesis and identify therapeutic strategies.

**Figure S1.**
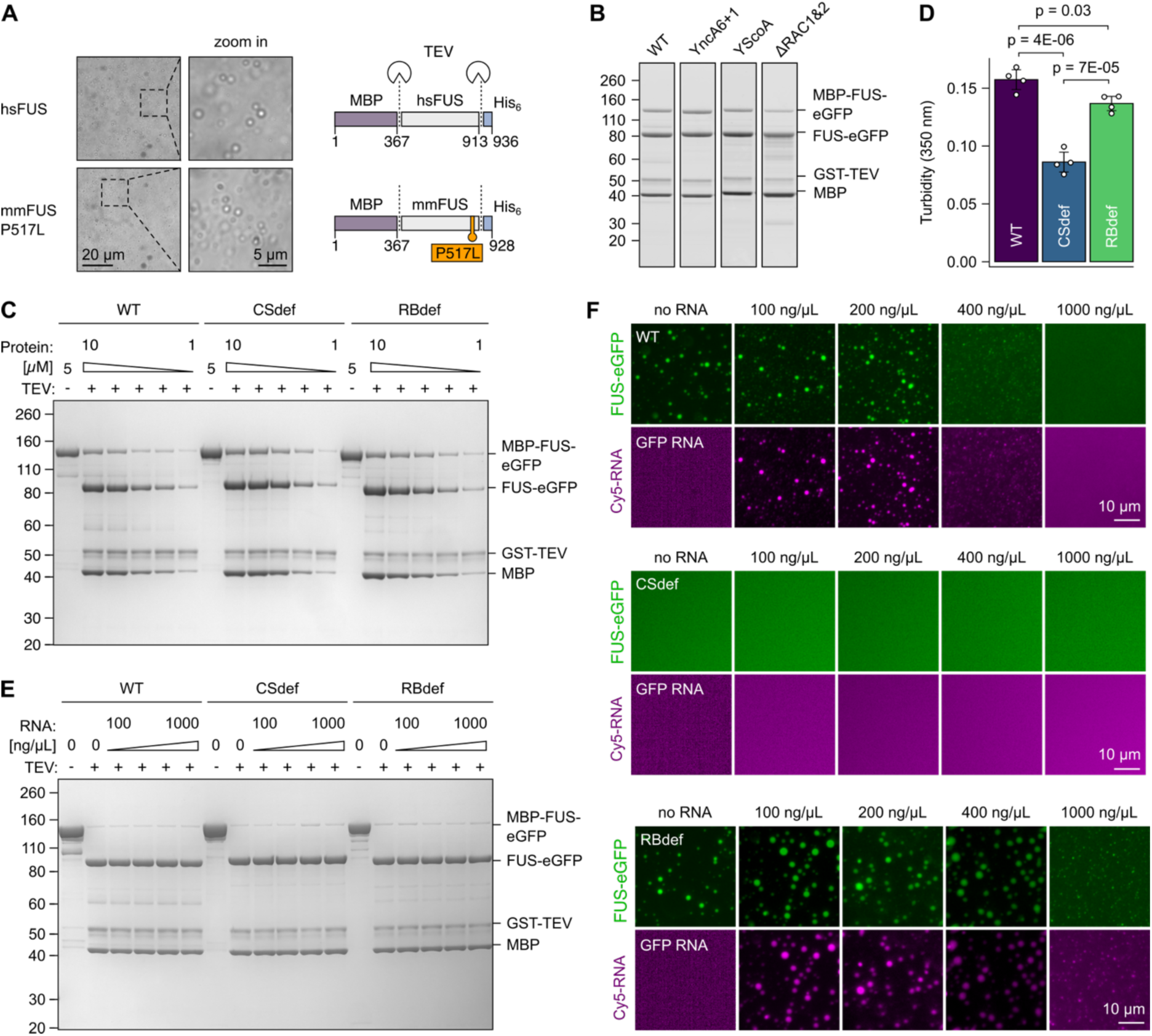
Identification of RNA-binding and condensation-deficient FUS mutants, related to Figure 1. (A) Brightfield microscopy images showing droplet formation of untagged human FUS (hsFUS) and mouse FUS carrying the P517L mutation (mmFUS-P517L) at 5 µM concentration, 30 min after TEV cleavage. Scale bar, 20 µm and 5 µm (zoom in). (B) Coomassie-stained SDS-PAGE gel confirming efficient cleavage of MBP-FUS-eGFP-His₆ constructs 30 min after TEV protease cleavage. (C) Uncropped Coomassie-stained SDS-PAGE gel corresponding to Figure 1E. (D) Turbidity at 350 nm used to quantify phase separation of 7 µM FUS-eGFP, 30 min after TEV cleavage. Data represent mean ± SD of N = 4 independent biological replicates. Statistical significance was determined by one-way ANOVA followed by Tukey’s Honest Significant Difference (HSD) test for multiple comparisons. (E) Coomassie-stained SDS-PAGE gel verifying efficient cleavage of MBP-FUS-eGFP-His₆ constructs in the presence of Cy5-labelled RNA. (F) Fluorescence microscopy images of WT, CSdef and RBdef FUS-eGFP (green) at 7.5 µM in the presence of increasing amounts of Cy5-labelled GFP RNA (magenta), 30 minutes post TEV-cleavage. Scale bar, 10 µm.

**Figure S2.**
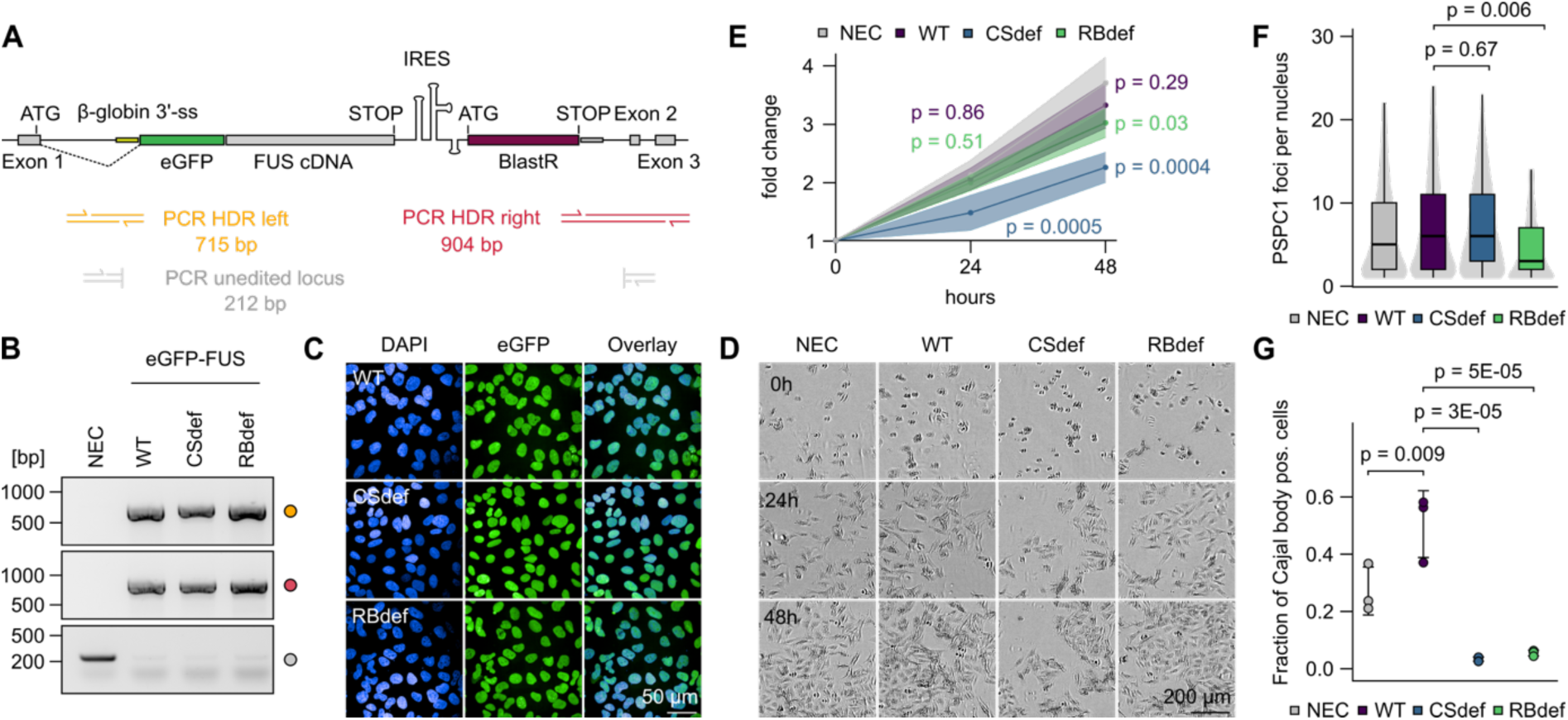
FUS condensation is required for paraspeckle recruitment and Cajal body formation, related to Figure 2. (A) Schematic depiction of the edited *FUS* locus and binding sites of primers used for genotyping by PCR across homology-directed repair (HDR) junctions. (B) Agarose gel confirming homozygous integration of the cDNA cassette into the endogenous *FUS* locus. (C) Fluorescence microscopy images of eGFP-FUS (green) and DAPI (blue) in edited U2OS cell lines showing purity of the cell lines and homogeneous expression. Scale bar, 50 µm. (D) Brightfield microscopy images of non-edited control (NEC) and edited U2OS cells at different timepoints after attachment. Scale bar, 200 µm. (E) Quantification of cell proliferation depicted as fold change relative to the initial timepoint. Data represent mean ± SD of N = 3 independent biological replicates. Statistical significance compared to non-edited control (NEC) was determined by one-way ANOVA for each timepoint, followed by Tukey’s Honest Significant Difference (HSD) test for multiple comparisons. (F) Violin-boxplot showing the number of PSPC1 foci per foci-positive nucleus. The plot displays median lines, interquartile range (IQR) boxes and 1.5 × IQR whiskers. n = 2,141 (NEC), 2,356 (WT), 2,808 (CSdef), 2,561 (RBdef) nuclei, from N = 3 independent biological replicates. Statistical significance was determined by one-way ANOVA followed by Tukey’s Honest Significant Difference (HSD) test for multiple comparisons. (G) Dot plot showing the fraction of cells containing Cajal bodies from Figure 2K. Data represent mean ± SD of N = 3 independent biological replicates. Statistical significance was determined by one-way ANOVA followed by Tukey’s Honest Significant Difference (HSD) test for multiple comparisons.

**Figure S3.**
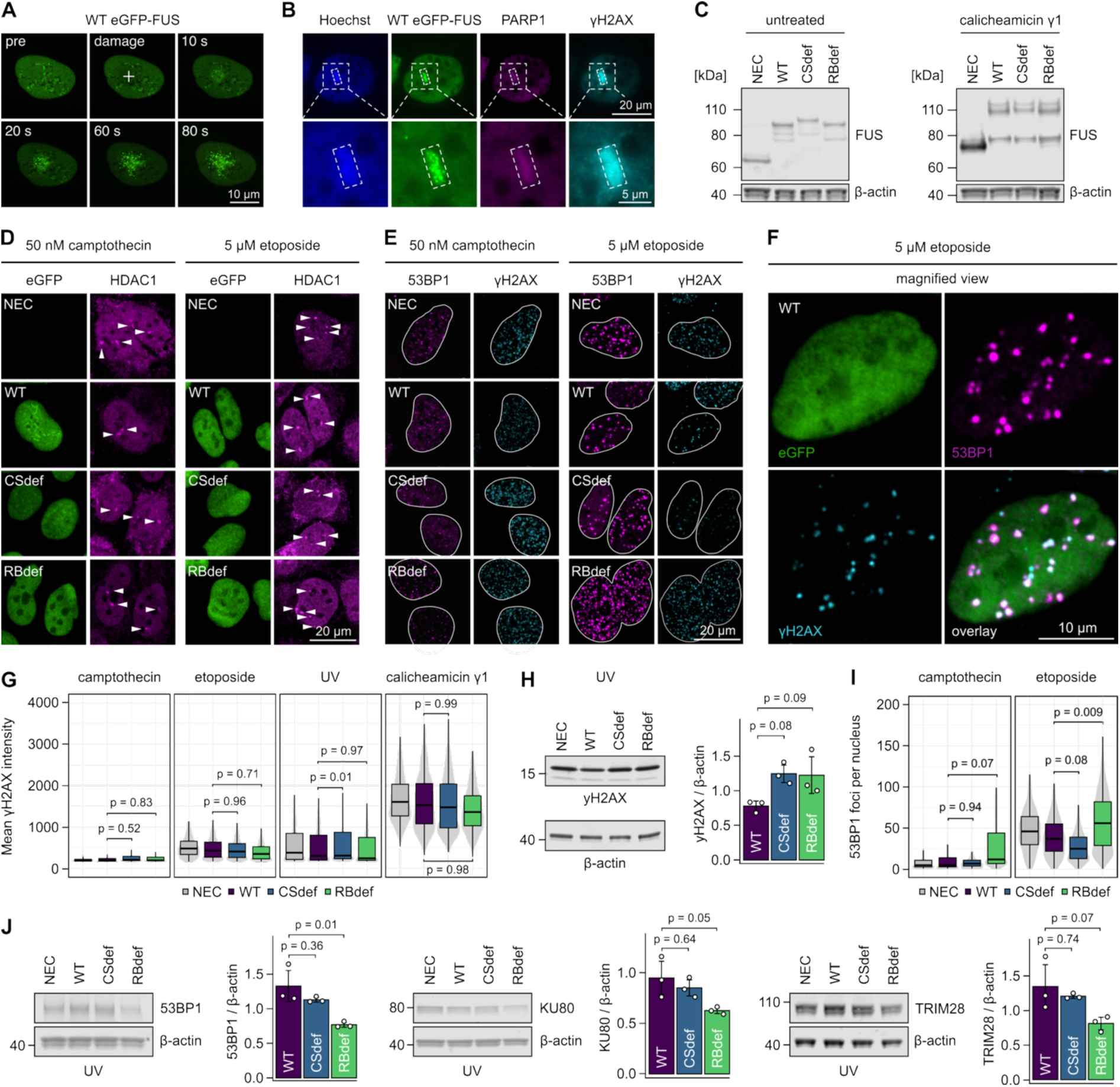
FUS condensation and RNA-binding distinctly contribute to FUS function in the DNA damage response, related to Figure 3. (A) Fluorescence microscopy images of live cells showing the localisation of eGFP-FUS before and at the indicated time points following laser-induced DNA damage. The point targeted by microirradiation is indicated by the white cross. Scale bar, 10 µm. (B) Fluorescence microscopy images of fixed cells showing the localisation of eGFP-FUS (green), PARP1 (magenta) and γH2AX (cyan) after DNA damage. The region of interest (ROI) targeted for microirradiation is outlined with dashed lines. Scale bar, 20 µm and 5 µm (zoom in). (C) Western blot analysis of FUS in untreated and calicheamicin-treated cells. The shifts to higher molecular weight indicate phosphorylation. (D) Fluorescence microscopy images showing the localisation of eGFP-FUS (green) and HDAC1 (magenta) upon induction of DNA damage with camptothecin and etoposide. White arrows indicate HDAC1 foci. Scale bar, 20 µm. (E) Fluorescence microscopy images of 53BP1 (magenta) and γH2AX (cyan) in camptothecin- and etoposide-treated cells. Nuclear outlines are defined by DAPI staining (white lines). Scale bar, 20 µm. (F) High magnification view of wild-type eGFP-FUS (green), 53BP1 (magenta) and γH2AX (cyan) upon etoposide treatment, from panel (E). Scale bar, 10 µm. (G) Violin-boxplot showing γH2AX intensity upon DNA damage induction. Camptothecin: n = 5,445 (NEC), 3,170 (WT), 3,737 (CSdef), 3,246 (RBdef); Etoposide: n = 1,784 (NEC), 1,818 (WT), 2,684 (CSdef), 2,345 (RBdef): UV: n = 1,224 (NEC), 1,397 (WT), 2,274 (CSdef), 1,783 (RBdef); Calicheamicin: n = 1,915 (NEC), 1,926 (WT), 2,853 (CSdef), 2,354 (RBdef) nuclei from N = 3 independent biological replicates. (H) Western blot analysis of γH2AX levels upon UV treatment. Bar plots show quantification relative to NEC. Data represent mean ± SD of N = 3 independent biological replicates. (I) Violin-boxplot showing the number of 53BP1 foci per positive nucleus upon DNA damage induction. Camptothecin: n = 4,695 (NEC), 2,367 (WT), 3,071 (CSdef), 3,094 (RBdef); Etoposide: n = 1,767 (NEC), 1,804 (WT), 2,660 (CSdef), 2,342 (RBdef) nuclei from N = 3 independent biological replicates. (J) Western blot analysis of 53BP1, KU80 and TRIM28 levels upon UV-treatment, with β-actin as loading control. Bar plots show quantification relative to NEC. Data represent mean ± SD of N = 3 independent biological replicates. All violin-boxplots (G, I) display median lines, interquartile range (IQR) boxes and 1.5 × IQR whiskers. All statistical tests (G, H, I, J) were performed by one-way ANOVA followed by Tukey’s Honest Significant Difference (HSD) test for multiple comparisons.

**Figure S4.**
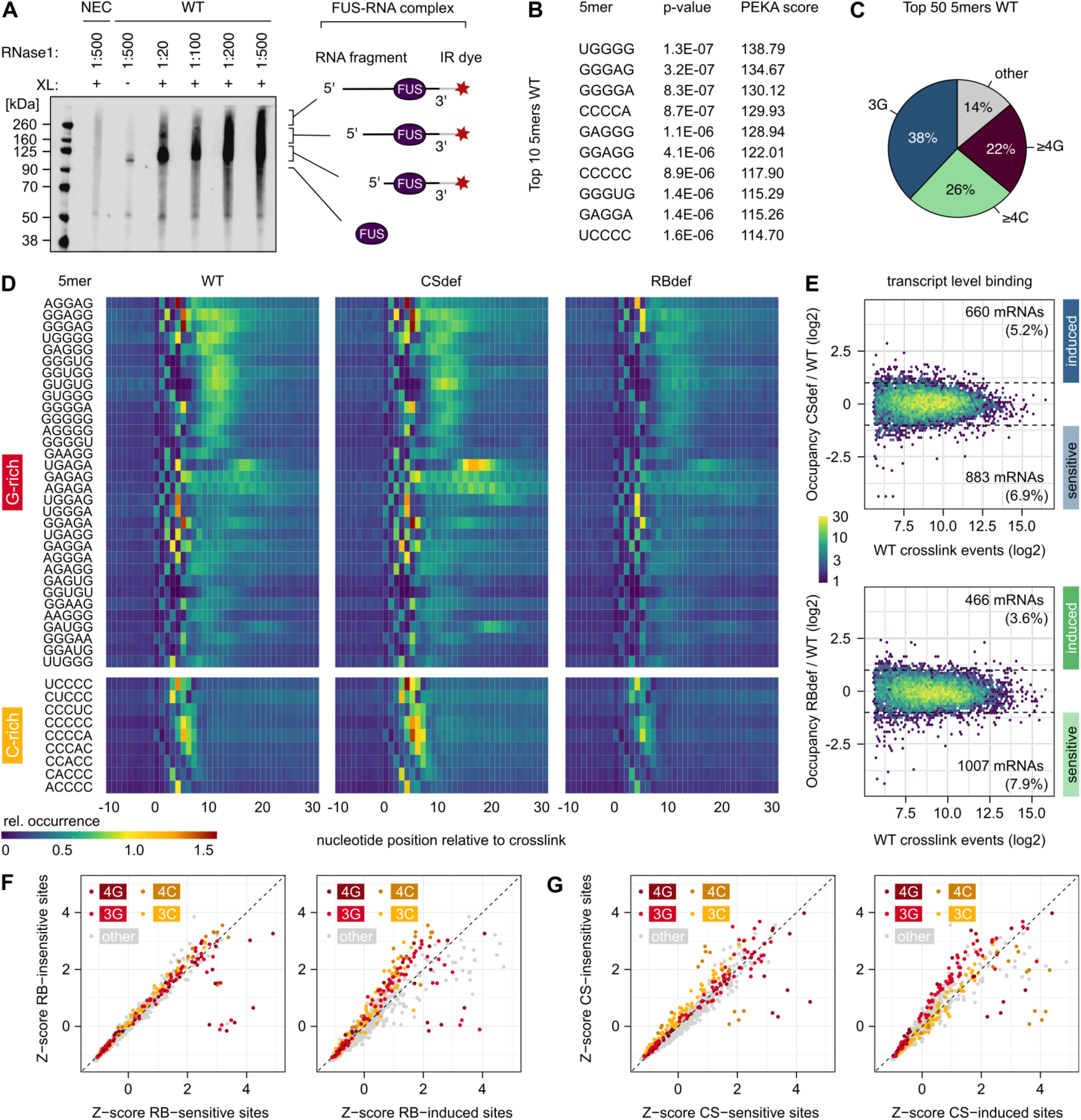
The RNA-binding landscape of FUS is shaped by condensation and canonical RNA-binding, related to Figure 4. (A) Visualisation of eGFP-FUS-RNA complexes following SDS-PAGE and transfer onto a nitrocellulose membrane. Non-edited control (NEC) cells and a no UV condition serve as specificity controls. Different RNase concentrations were tested to optimise the length of the cross-linked RNA fragments. (B) Top 10 significantly enriched 5-mers identified in wild-type eGFP-FUS crosslink clusters. (C) Sequence composition of the 50 most enriched 5-mers. Pie chart shows the fraction of top 50 5-mers containing ≥ 4 guanosines (4G, dark red), 3 guanosines (3G, blue), ≥ 4 cytosines (4C, green). The remaining 5-mers are shown in grey. (D) Heatmaps showing the positional enrichment of G-rich and C-rich 5-mers (from the top 50 motifs) relative to the crosslink sites in wild-type, CSdef and RBdef eGFP-FUS clusters. (E) Hexagonal binning plots showing transcript-level eGFP-FUS occupancy of CSdef (top) and RBdef (bottom) eGFP-FUS relative to WT, plotted against total WT crosslink events for each transcript. Only protein-coding transcripts not differentially expressed between cell lines are included. The colour scale indicates point density. Dashed lines indicate two-fold changes in either direction. (F) Scatterplots comparing 5-mer frequency (Z-scores) between RNA-binding-sensitive and -insensitive (left) or loss-of-RNA-binding-induced and -insensitive (right) FUS binding sites. 5-mers containing ≥ 4 guanosines (4G, dark red), 3 guanosines (3G, red), ≥ 4 cytosines (4C, dark orange), 3 cytosines (3C, orange) are highlighted. (G) Scatterplots comparing 5-mer frequency (displayed as Z-scores) between condensation-sensitive and -insensitive (left) or loss-of-condensation-induced and -insensitive (right) FUS binding sites. G-rich and C-rich 5-mers are coloured as in panel (F).

**Figure S5.**
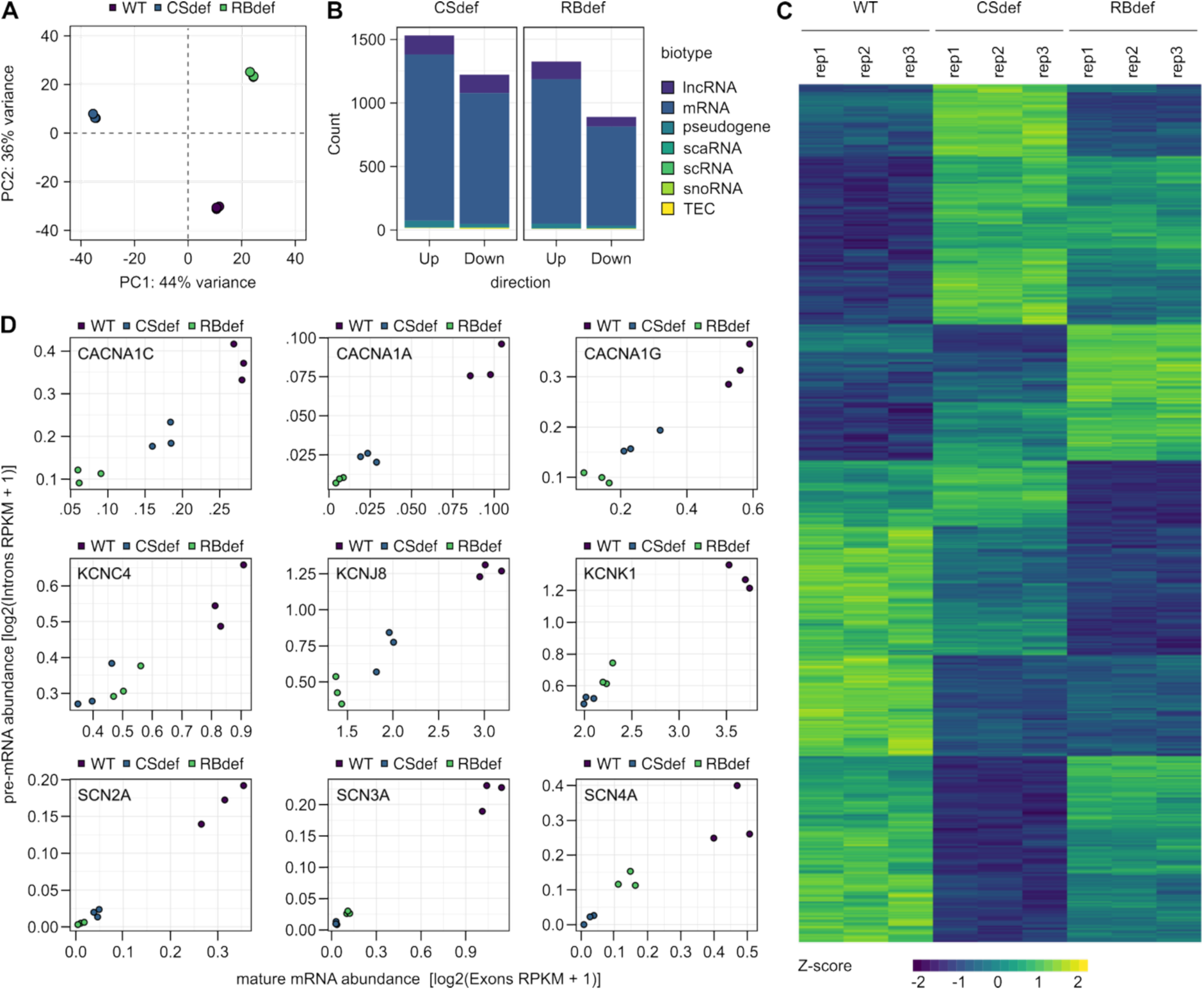
FUS condensation and RNA-binding contribute to ion channel mRNA expression, related to Figure 5. (A) Principal component analysis showing the variance in gene expression across the different U2OS cell lines. Each point represents one biological replicate. Clustering of replicates within each cell line indicates high reproducibility in their expression profiles. (B) Stacked bar plots summarising the number of significantly up- and downregulated genes (Fold-change > 1.5 and adjusted p-value < 10E-05) in CSdef and RBdef cells compared to wild-type, according to biotype. Protein-coding genes are the largest category, followed by long non-coding RNAs (lncRNAs). (C) Heatmap showing relative expression levels of differentially expressed genes (Fold-change > 1.5 and adjusted p-value < 10E-05) across cell lines and biological replicates. Each row represents a gene. Expression values represent SD from mean of variance stabilised values across rows. Hierarchical clustering reveals both similarities and differences in gene expression patterns between the cell lines. (D) Dot plots comparing the abundance of mature mRNA (based on exonic reads) and pre-mRNA (based on intronic reads) for selected ion channel mRNAs. Expression values are shown as log2(Reads Per Kilobase of transcript per Million mapped reads (RPKM) + 1). The strong correlation between exonic and intronic signals indicates transcriptional regulation of these genes.

**Figure S6.**
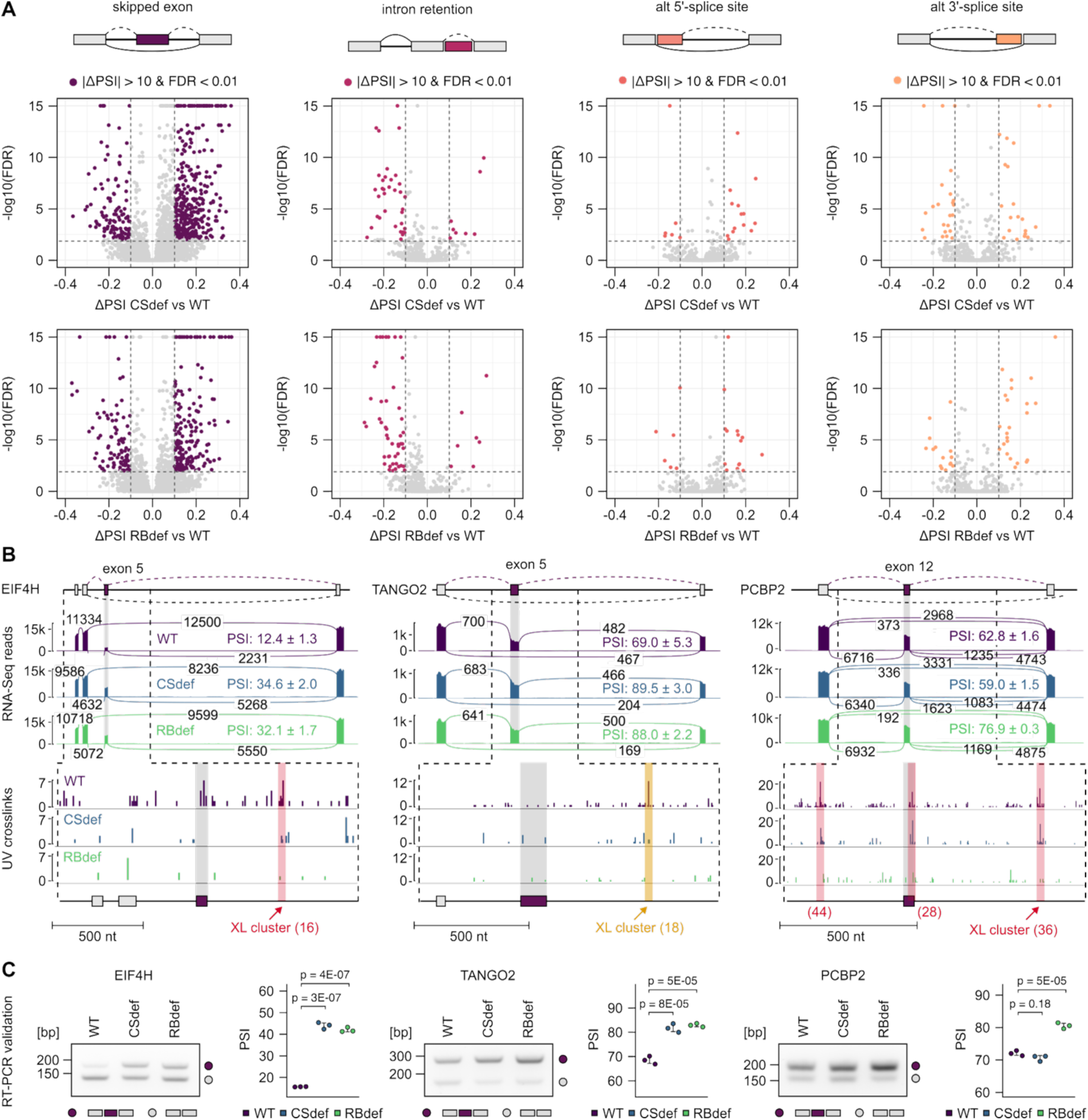
FUS condensation and RNA-binding co-operatively regulate alternative splicing, related to Figure 6. (A) Volcano plots showing differences in alternative splicing between CSdef (top) and RBdef (bottom) eGFP-FUS cells lines compared to wild-type, separated by type of alternative splicing. Vertical dashed lines indicate an absolute change in percent spliced in (|ΔPSI|) > 10, horizontal lines mark the significance threshold (false discovery rate (FDR) < 0.01). (B) Genome browser view of *EIF4H, TANGO2* and *PCBP2* regions containing FUS-regulated alternative exons. Top: Sashimi plots showing RNA-Seq junction reads and PSI values. Bottom: Bar plots of eGFP-FUS crosslinks around regulated exons, with G-rich (red) and C-rich (orange) clusters highlighted. (C) RT-PCR validation of alternative splicing in *EIF4H, TANGO2* and *PCBP2*. Dot plots show PSI values (Mean ± SD of N = 3 independent biological replicates). Statistical significance was analysed by one-way ANOVA followed by Tukey’s HSD test for multiple comparisons.

## Resource availability

### Lead contact

Further information and requests for resources and reagents should be directed to the lead contact, Marc-David Ruepp.

### Materials availability

Plasmids and cell lines generated in this study are available from the lead contact.

## Supporting information

TableS1 - DESeq2 differential expression analysis

TableS2 - rMATS analysis SE

TableS3 - rMATS analysis RI

TableS4 - rMATS analysis A5SS

TableS5 - rMATS analysis A3SS

TableS6 - Recombinant DNA constructs

TableS7 - Oligonucleotide sequences

TableS8 - Antibodies

## Acknowledgements

We thank George Chennel and Chen Liang from the Wohl Cellular Imaging Centre, (King’s College London, London, UK) for training and technical support with microscopy techniques. We thank Jernej Ule (King’s College London, London, UK) for stimulating discussions and technical advice on this work. This research and related results were made possible through the support of the NOMIS Foundation [MD.R. & D.D.] and the UK Dementia Research Institute [award number UK DRI-6204 to MD.R.] through UK DRI Ltd, principally funded by the Medical Research Council. J.A. is supported by a Motor Neurone Disease Association (MNDA) PhD studentship (Ruepp/Oct22/911-792) awarded to MD.R. and H.PF. D.J. is supported by an Alzheimer’s Research UK fellowship (ARUK-RF2024-002). C.R.S. is supported by a Sir Henry Dale Fellowship, jointly funded by the Wellcome Trust and the Royal Society (215454/Z/19/Z). This research was also supported by the Deutsche Forschungsgemeinschaft (DFG) (Heisenberg grant, project number 44269835 and RTG 2859 “R-loop regulation in robustness and resilience”, project number 491145305) to D.D.

## Author contributions

Conceptualisation, MD.R. & D.D.; methodology, MD.R., D.D., D.J., J.A., F.T., C.R.S.; investigation, D.J., J.A. & S.H.; resources, B.D.; visualisation, D.J., J.A.; funding acquisition: C.R.S., H.PF., D.D. & MD.R.; project administration, MD.R., D.D & D.J.; supervision, C.R.S., H.PF., D.D & MD.R.; writing - original draft, D.J.; writing - review & editing, D.J., J.A., MD.R.; writing - final editing/approval of final version, all authors.

## Declaration of generative AI and AI-assisted technologies in the writing process

During the preparation of this manuscript, the authors used ChatGPT and Grammarly to assist with language refinement and to enhance readability. The authors take full responsibility for the content of the article.

## STAR Methods

### Key resources

A complete list of recombinant DNA constructs (Table S6), oligonucleotides (Table S7), and antibodies (Table S8) used in this study is provided.

### Experimental model and study participant details Cell culture

U2OS cells (ECACC, Acc N° 92022711) were maintained at 37°C and 5% CO_2_, in DMEM/F12 GlutaMAX (Gibco, #31331028) supplemented with 10% (v/v) fetal bovine serum (FBS) (PAN-Biotech, #P30-3031) and 100 U/ml (1% v/v) Penicillin-Streptomycin (Gibco, #15140122), and passaged using TrypLE Express (Gibco, #12605010) unless stated differently.

### U2OS genome-editing

Genome editing of all cell lines was done following our previously published CRISPR-Trap strategy^46^. In brief, cells were transfected with a plasmid expressing Cas9 and a guide RNA targeting *FUS* intron 1 (plasmid ID: I20), and plasmids expressing homology direct repair (HDR) gene replacement matrices encoding the different eGFP-FUS variants (plasmid IDs: I123-125), in a molar ratio of 1:1.17 using *Trans*IT-LT1 Transfection Reagent (Mirus Bio, #MIR 2300) in Opti-MEM Reduced Serum Medium (Gibco). 48h after transfection, cells were selected with 10 µg/ml blasticidin S HCl (Gibco, #A1113903) for 5 days. After antibiotic selection, cells were dissociated and seeded as single cells in 15 cm cell culture dishes. eGFP-positive colonies were isolated using cloning cylinders and transferred to single wells of 24-well plates for expansion. Clones were screened by WB using α-FUS and α-GFP antibodies to verify expression of eGFP-FUS and loss of endogenous FUS protein. Potential homozygous clones were further validated by PCR genotyping with APA Taq EXtra HotStart ReadyMix (Roche, #KK3605) using primers spanning the left HDR junction (primer IDs: SRE198/JA70), the right HDR junction (primer IDs: JA81/MDR352) as well as primers spanning the *FUS* gRNA target region (primer IDs: JA59/SRE199). Genomic DNA (gDNA) was extracted using the DNeasy Blood & Tissue kit (Qiagen, #69504) or QuickExtract DNA Extraction Solution (Biosearch Technologies, #QE09050) following manufacturer’s instructions.

### Method details Plasmid cloning

Eukaryotic expression constructs were generated as follows: pcDNA3.1(+)-FLAG-mmFUS-WT (Plasmid ID: B127), pcDNA3.1(+)-FLAG-mmFUS-RBdef (Plasmid ID: B128) and pcDNA3.1(+)-FLAG-mmFUS-P517L (Plasmid ID: B130) were generated by gene synthesis (GeneArt, Thermo). To create the YncA6+1, YScoA and ΔRAC1&2 variants (Plasmid IDs: B137, B138 and B139), gene-synthesised cDNA encoding mouse FUS amino acids 1 - 252 with the desired mutations was inserted into the HindIII and SacII sites of pcDNA3.1(+)-FLAG-mmFUS-P517L (Plasmid ID: B130). To create pcDNA6-eGFP-GSG15-mmFUS-WT (C77) and pcDNA6-eGFP-GSG15-mmFUS-RBdef (C79), the entire FUS coding regions of pcDNA3.1(+)-FLAG-mmFUS-WT (Plasmid ID: B127) and pcDNA3.1(+)-FLAG-mmFUS-RBdef (Plasmid ID: B128) were PCR amplified (Primer IDs: JA8 and JA9) and inserted into the XhoI and NotI sites of pcDNA6-eGFP-GSG15-hsFUS-WT (Plasmid ID: C19)^77^. To create pcDNA6-eGFP-GSG15-mmFUS-YncA6+1 (Plasmid ID: C80), gene-synthesised cDNA encoding mouse FUS amino acids 1 - 252 with the desired mutations was PCR amplified (Primer IDs: JA8 and DJ790) and inserted into the XhoI and SacII sites of pcDNA6-eGFP-GSG15-mmFUS-WT (C77).

Bacterial expression constructs were generated as follows: pMal-C5-TEV-TEV-His (Plasmid ID: A61) and pMal-C5-TEV-eGFP-TEV-His (Plasmid ID: A62) were generated by cloning of the respective gene-synthesised fragments (General Biosystems) into the SacI and HindIII sites of pMal-C5, with in-frame XbaI and BamHI sites downstream of the first TEV cleavage site, and upstream of TEV-His or eGFP-TEV-His, respectively. To generate pMal-C5-TEV-mmFUS-P517L-TEV-His (Plasmid ID: G26), the FUS coding region of pcDNA3.1(+)-FLAG-mmFUS-P517L (Plasmid ID: B130) was PCR amplified (Primer IDs: MDR948 and MDR950) and inserted into the XbaI and BamHI sites of pMal-C5-TEV-TEV-His (Plasmid ID: A61). To generate pMal-C5-TEV-mmFUS-P517L-eGFP-TEV-His (Plasmid ID: G34) and the YncA6+1, YScoA, ΔRAC1&2 and RBdef variants (Plasmid IDs: G37, G38, G39 and G40), the FUS coding region of the corresponding pcDNA3.1 constructs (Plasmid IDs: B130, B137, B138, B139 and B128) was PCR amplified (Primer IDs: MDR948 and MDR950) and inserted into the XbaI and BamHI sites of pMal-C5-TEV-eGFP-TEV-His (Plasmid ID: A62).

Gene editing constructs were generated as follows: pMK-HDR-FUSKI-G515Vfs-Puro (Plasmid ID: I110) and a BSD-SV40pA-HDR string were generated via synthesis (GeneArt, Thermo). The BSD-SV40pA-HDR string was then cloned into BmgBI and PacI sites of pMK-HDR-FUSKI-G515Vfs-Puro (Plasmid ID: I110), resulting in pMK-HDR-FUSKI-G515Vfs-BSD (Plasmid ID: I115). To create pMK-HDR-eGFP-mmFUS-WT-BSD (Plasmid ID: I123), pMK-HDR-eGFP-mmFUS-YncA6+1-BSD (Plasmid ID: I124) and pMK-HDR-eGFP-mmFUS-RBdef-BSD (Plasmid ID: I125), the β-globin chimeric intron region was amplified (Primer IDs: DJ852 and DJ853) from pMK-HDR-FUSKI-G515Vfs-BSD (Plasmid ID: I115). The eGFP-mmFUS coding region was amplified (Primer IDs: DJ854 and JA9) from pcDNA6-eGFP-GSG15-mmFUS-WT (Plasmid ID: C77), pcDNA6-eGFP-GSG15-mmFUS-YncA6+1 (Plasmid ID: C80) and pcDNA6-eGFP-GSG15-mmFUS-RBdef (Plasmid ID: C79). The two fragments were then fused by PCR (Primer IDs: DJ852 and JA9) and cloned into the BamHI and NotI sites of pMK-HDR-FUSKI-G515Vfs-BSD (Plasmid ID: I115). pU6gRNA-Cas9-RFP (Plasmid ID: I20) targeting *FUS* intron 1 (target sequence TGGATGTCCACCAAGACCTTGG) according to^46^ was generated by gene synthesis (Sigma Aldrich).

### Recombinant protein expression and purification

Recombinant mouse FUS protein was expressed and purified using a modified version of the protocol described in^28^. Expression plasmids encoding MBP-FUS-eGFP-His proteins were transformed into chemically competent *E. coli* Rosetta-2(DE3)-pLysS cells (Novagen, #714014) and grown in 2 L lysogeny broth (LB) medium containing carbenicillin (100 µg/mL) and chloramphenicol (33 µg/mL). At an OD_600_ of 0.6 - 0.8, protein expression was induced with 0.1 mM IPTG for 24 hours at 12°C. Bacteria were harvested by centrifugation at 4,000 × g for 20 minutes and pellets were stored at -80°C. Frozen pellets were resuspended in 10 mL lysis buffer (300 mM NaCl, 20 mM Imidazole, 20 µM ZnCl_2_, 10% glycerol in 50 mM NaH_2_PO_4_ pH 8.0) supplemented with 4 mM β-mercaptoethanol, EDTA-free protease inhibitor (Roche, #11836170001), RNase A (Sigma, #10109169001), and DNase I (Roche, #04716728001). After sonication, lysates were cleared by centrifugation at 24,500 × g for 1 hour at 10°C, and the supernatant was filtered through a 0.2 µm PES membrane. The cleared lysate was subjected to affinity purification using Ni-NTA agarose resin (Qiagen, #30410). For each construct, a total of 15 mL 50% slurry (7.5 mL resin) was equilibrated in lysis buffer and incubated with the lysate for 2 hours at 4°C on a rotator. After batch binding, the resin was transferred to a gravity flow column (BioRad, #7372512), washed with 100 mL lysis buffer, and eluted in five 5 mL fractions using His-Trap elution buffer (300 mM NaCl, 250 mM Imidazole, 20 µM ZnCl_2_ in 50 mM NaH_2_PO_4_ pH 8.0). Aliquots of each elution were analysed by SDS-PAGE. Selected fractions were pooled for further purification. For each construct, a total of 15 mL 50% slurry (7.5 mL resin) Amylose resin (NEB, #E8021) was equilibrated in wash buffer (300 mM NaCl, 20 mM imidazole, 20 µM ZnCl_2_ in 50 mM NaH_2_PO_4_ pH 8.0) and combined with the pooled Ni-NTA eluates in a 50 mL tube. The mixture was incubated overnight at 4°C with gentle rotation. On the following day, ionic strength was adjusted to 150 mM NaCl by diluting the resin 1:1 with salt-free wash buffer. After an additional 1-hour incubation at 4°C, the beads were transferred to a gravity column, washed with 50 mL wash buffer, and the protein was eluted in 5 mL fractions using amylose elution buffer (wash buffer with 20 mM maltose). Elution fractions were analysed by SDS-PAGE. High-yield fractions were pooled and concentrated using 100 kDa molecular weight cutoff centrifugal filters (Sartorius, #VS2042) at 3,000 × g and 4°C until reaching a final concentration of approximately 4 mg/ml. Protein concentration was measured using a Nanodrop spectrophotometer with a calculated extinction coefficient of 161.36 mM⁻¹cm⁻¹ and a molecular weight of 125.7 kDa. Purified proteins had 260/280 nm ratios between 0.6 and 0.8. Glycerol was added to a final concentration of 5%, and the purified protein was aliquoted, flash-frozen in liquid nitrogen, and stored at -80°C until use. Prior to all *in vitro* phase separation assays, purified MBP-FUS-eGFP-His proteins were thawed on ice and centrifuged at 21,000 x g for 20 min at 4°C to remove any insoluble aggregates. Salt concentration was then adjusted to 150 mM NaCl by diluting the protein 1:1 with salt-free droplet buffer (20 µM ZnCl_2_ in 50 mM NaH_2_PO_4_ pH 8.0).

### *In vitro* transcription

GFP mRNAs (774 nt) used in phase separation assays was *in vitro* transcribed from the T7 promoter of pcDNA3.1-FLAG-eGFP (Plasmid ID: B26) linearised with XbaI (NEB, #R0145). Transcription was performed using the MEGAshortscript T7 kit(Thermo Fisher Scientific, #AM1354) in the presence of 10 mM Cy5-UTP (APExBio, #B8333) to fluorescently label the transcripts. Reactions were incubated at 37°C for 4 hours, followed by digestion of template DNA with Turbo DNase for 30 minutes at 37°C. Transcribed RNA was purified by phenol:chloroform extraction and ethanol precipitation. RNA concentration was quantified using a Nanodrop spectrophotometer, and transcript integrity was verified by agarose gel electrophoresis.

### Droplet assay

For droplet assays, the proteins and *in vitro* transcribed RNAs were diluted to the desired concentrations in droplet buffer (150 mM NaCl, 20 µM ZnCl_2_ in 50 mM NaH_2_PO_4_ pH 8.0) and phase separation was induced upon addition of TEV protease (Sigma, #T4455). Half of the reaction was immediately transferred to an 18-well chamber slide (Ibidi, #81816), where droplet formation was visualised with a Nikon Ti-E eclipse epifluorescence microscope and NIS elements AR imaging software (Nikon, v5.01), 30 minutes after TEV cleavage. The remaining half of the reaction was used to confirm TEV cleavage and integrity of the proteins at the time of imaging. Hereto, the proteins were mixed with LDS sample buffer 30 minutes post TEV cleavage, followed by analysis via SDS-PAGE and Coomassie staining.

### Sedimentation assay

For sedimentation assays, the proteins were incubated with TEV protease in droplet buffer for 60 minutes and then centrifuged at 21,000 x g for 15 minutes to pellet phase-separated FUS. After transfer of the supernatants to fresh tubes, the pellets were dissolved in an equal volume of droplet buffer. The samples were then separated on 4-12% polyacrylamide gels, fixed for 30 minutes in 50% EtOH, 10% acetic acid, and stained with SyPro Ruby (Invitrogen, #S12000) overnight. Following destaining in 10% EtOH, 10% acetic acid for 30 minutes, the gels were visualised on a Typhoon FLA 9000 Fluorescent scanner (Amersham, GE healthcare). Quantitative measurements were performed using Fiji (v2.0.0)^78^.

### Turbidity assay

For turbidity measurements, proteins were diluted to 7 µM in turbidity buffer: 150 mM NaCl, 2 mM DTT, 20 mM NaH_2_PO_4_ pH 8.0. After 30 minutes of TEV protease cleavage, the reactions were transferred in a 384-well plate, and turbidity was measured at 600 nm and 350 nm on a SpectraMax ID5 plate reader (Molecular Devices). For each biological replicate, two technical replicates were recorded, and background absorbance (buffer + TEV) was subtracted.

### RT-PCR and RT-qPCR

RNA was isolated using the Absolutely RNA miniprep kit (Agilent, #400805) and reverse transcribed into cDNA using LunaScript RT SuperMix (NEB, #E3010) following the manufacturer’s protocols. RT-PCR was carried out for 25 cycles using GoTaq Green Master Mix (Promega, #M712) in 30 µl reactions containing 32 ng cDNA and 666 nM primers each. qPCR was performed using PowerUp SYBR Green Master Mix (Thermo, #A25742) in 20 µl reactions containing 32 ng cDNA and 600 nM primers each. Amplification and fluorescence monitoring was performed on a CFX96 Real-Time PCR System (BioRad) using CFX maestro software (v2.2). The standard two-step cycling program recommended by the manufacturer (Thermo) was employed, followed by a melting curve analysis to verify product specificity. Relative transcript levels were computed using the ΔΔCt method^79^.

### Western blot

Western blots (WB) were carried out following standard protocols. Generally, cells were lysed in RIPA buffer (Thermo Scientific, #89901) supplemented with Halt Protease Inhibitor Cocktail (Thermo Scientific, #87786) on ice for 15 min. After clearance of cellular debris by centrifugation at 16,000 x g for 5 min, the RIPA extracted supernatants were combined with NuPAGE 2x LDS Sample Buffer (Invitrogen, #NP0008) containing 200 mM dithiothreitol (DTT, PanReac, #A3668) and heated at 95°C for 5 min to denature proteins prior to sodium dodecyl sulphate-polyacrylamide gel electrophoresis (SDS-PAGE). For the reliable detection of post-translationally modified or chromatin-associated proteins, RIPA buffer was supplemented with phosphatase inhibitor (Thermo Scientific, #181280), 1M MnSO_4_ (Sigma, #M8179), 500 U/ml benzonase (Sigma, #E1014) and 50 µg/ml RNase (Sigma, #R6148), and the debris clearance step was omitted. SDS-PAGE was performed using 4-12% or 12% NuPAGE Bis-Tris or 3-8% Tris-Acetate gradient protein gels (Invitrogen), in NuPAGE MOPS (Invitrogen, #NP000102) or Tris-Acetate (Invitrogen, #LA0041) SDS running buffers. Separated proteins were transferred to nitrocellulose membranes using the iBlot 2 transfer system (Invitrogen) for 7 min at 20V or for 10 min at 20V for high-molecular-weight proteins (e.g., 53BP1). To confirm efficient transfer and as a secondary loading control, membranes were stained with Ponceau S (Abcam, #ab270042) for 1 min, followed by de-staining with deionised water. Then, membranes were blocked 1h at RT with 5% (w/v) skimmed milk (Serva, #42590.02) in TBS-T (TBS, 0.1% (v/v) Tween, 0.1% (w/v) NaN_3_) (Sigma, #71290) and incubated with primary antibodies overnight at 4°C. For the detection of phosphorylated proteins, a 5% (w/v) bovine serum albumin (BSA, Sigma, #A9418) solution in TBS-T was used instead of milk. Following washes with TBS-T, membranes were incubated with infrared dye-conjugated secondary antibodies for 1 hour at RT. Then, membranes were washed, scanned using an Odyssey CLx imaging system (Li-Cor) and processed with Image Studio software (Li-Cor, v5.2). Quantitative measurements were performed using Fiji (v2.0.0)^78^.

### Immunocytochemistry (ICC)

ICC was generally performed as follows: Cells were washed with PBS and fixed with a PFA 4% solution (Thermo Fisher, #J19943.K2) at RT for 15 min. After washing with PBS, the cells were blocked and permeabilised in 1x TBS, 0.5% (v/v) Triton-X-100 (PanReac, #A4975,0500), 6% (w/v) bovine serum albumin (BSA, Sigma, #A9418) for 30 min. Primary antibodies were diluted in antibody dilution buffer (1x TBS, 0.1% (v/v) Triton-X-100, 6% (w/v) BSA) and incubated overnight at 4°C. The next day, cells were washed three times 10 minutes with PBS and incubated with Alexa Fluor-conjugated secondary antibodies and 4′,6-diamidino-2-fenilindol (DAPI, Life Technologies, #834650) for 1h at RT. Following another three 10-minute washes with PBS, the cells were kept at 4°C in PBS until imaging. Image acquisition was done using an Opera Phenix Plus (Perkin Elmer) or an ECHO Revolve fluorescence microscope (BICO). For printing, brightness and contrast of the pictures were linearly enhanced.

### Cell proliferation

Cells were seeded in PhenoPlate cell imaging microplates (Revvity, #6055302) and incubated at 37°C and 5% CO_2_ inside an IncuCyte SX5 live-cell analysis system (Sartorius). 9 brightfield pictures (fields) per well were taken in the same position using a 20× lens 4h after cell seeding, recovery and attachment (0h), and at 24h and 48h after the first timepoint. Total cell number count per image/field was automatically analysed and calculated using CellProfiler (Broad Institute, v4.2.8)^80^. Alternatively, in order to confirm the growth trends in later timepoints avoiding confluency issues for automated analysis in brightfield, the same cell number was simultaneously seeded in five PhenoPlate microplates, fixed with 8% PFA (1:1, v/v) (Thermo Scientific, #047347.9M) 4h after seeding (0h) and every 24h after initial timepoint up to 96h, and stained with DAPI for 15 min at room temperature (RT). 97 pictures (fields) per well were taken with a 20× air lens in the DAPI channel using an Opera Phenix Plus high-content imaging system (Perkin Elmer), and automated nuclei count was done using Harmony software (Perkin Elmer, v4.9.2137.273).

### Laser microirradiation assay

Glass-bottom culture dishes (Thermo Scientific, #150680) were coated for 1h at RT with poly-d-lysine (Gibco, #A3890401) and washed with warm media before cell seeding to remove any residual coating agent. Cells were seeded and incubated overnight for proper settling, attachment and recovery. Before laser microirradiation, cells were pre-sensitised for 30 min with 0.5 µg/ml bisBenzimide H 33342 trihydrochloride (Hoechst, Sigma, #B2261) in DMEM Fluorobrite (Gibco, #A1896701) supplemented with 2% FBS. After incubation with Hoechst, cells were used for the experiment only up to a maximum of 30 min to avoid viability issues. Laser microirradiation was performed at 37°C and in a 5% CO_2_ atmosphere in a Nikon AX inverted confocal microscope equipped with an Okolab incubator with CO_2_ control and using a 40× water immersion lens (Apo LWD 40x WI λS DIC N2). Three images were taken as baseline (pre-irradiation), then cells were microirradiated within a rectangular ROI for 1s using the 405 nm laser at 5% power, and eGFP-FUS recruitment was recorded up to 4 min using NIS-Elements AR software (Nikon, v5.42.05). Experiments were carried out in triplicate on three different days, including technical replicas for each cell line on each day. Acquisition parameters: 488 nm channel, Frame size 512 x 512 pixels (37.44 x 37.44 µm), Averaging: 2, Galvano bidirectional scanning, Scan mode: band, Dwell time: 1 µsec (1 fps), detector: 2, gain: 20, Pinhole: 22.4 µm, Scan area: zoom size = 11.8, Nyquist = 0.102 µm. Microirradiation parameters: 5% laser power on 405 nm laser, 1 iteration of 1s. Microirradiation area: rectangle of 5.4 x 0.18 µm. Acquisition cycles: 240 cycles, frame interval = 969.7 ms. Analysis was performed with FIJI (v2.0.0) as described in^81^. First, the fluorescence intensities within the microirradiated ROIs as well as size-matched control ROIs in the same nuclei at each time point were normalised to the corresponding intensities pre-irradiation. To account for the bleaching due to prolonged imaging, the baseline-normalised intensity values for the control ROIs were then subtracted from the irradiated ROIs and converted into percentages for data visualisation.

### UV- and chemical-induced DNA damage assays

Cells were seeded in PhenoPlate cell imaging microplates, incubated overnight at 37°C and, where indicated, treated for 1h with 50 nM camptothecin (CPT, #CAY11694), 5 µM etoposide (ETO, #CAY12092) or 50 nM calicheamicin γ1 (CAL, #CAY40830). A final 0.025% (v/v) DMSO (Apollo Scientific, #67-68-5) was added to non-treated controls. For UV-induced DNA damage, cells were irradiated in PBS with 8 mJ/cm^2^ UV-C (254 nm) inside a CL-1000 UVP Crosslinker (Analytik Jena) and recovered in media at 37°C for 4h before fixing for ICC processing. For western blotting, cells were seeded in 60 mm cell culture dishes and treated as outlined above.

### High-throughput image acquisition and analysis

Imaging of all eGFP-FUS foci, paraspeckles, Cajal bodies and DNA-damage repair foci was carried out using an Opera Phenix Plus high-content screening system (Perkin Elmer) using 40× or 60× water-immersion lenses in confocal mode. Z-stacks were taken every 0.5 µm covering the whole 3D area of cells. Acquisition for each channel was kept separated through all scans to avoid crosstalk between different fluorescent signals. Quantitative analysis of the different readouts was performed using the maximum projection of all Z-stacks using Harmony software (Perkin Elmer, v4.9.2137.273).

### Electrophoretic mobility shift assay (EMSA)

The recombinant FUS-RBD constructs were already described in^43^. Cy5-labelled RNA oligonucleotides were ordered from IDT. Stock solutions (100 µM) were diluted to 200 nM in 1x EMSA binding buffer: 50 mM KCl (Invitrogen, #AM9640G), 10 µg/ml yeast tRNA (Invitrogen, #AM7119), 50 µg/ml BSA (Sigma, #A3059), 10 mM HEPES pH 7.3 (Severn Biotech, #20-7200-01). For the binding reactions, 1.6 pmol RNA (80 nM final concentration) were mixed with serially diluted FUS-RBD constructs (8 nM to 2 µm final concentration) in 1x EMSA binding buffer and incubated at RT for 1 hour. Then, RNA gel loading buffer, 5% glycerol, traces of Orange G (Sigma, #O3756), was added and protein-RNA complexes were separated by non-denaturing electrophoresis in a 6% polyacrylamide (29:1, Sigma, #A7802) gel in 0.5x TBE buffer (Invitrogen, #15581-044) on ice. The gels were subsequently imaged on an Odyssey Fc imaging system (Li-Cor) with Image Studio software (Li-Cor, v5.2)

### RNA sequencing

We performed total RNA sequencing on three biological replicates each of WT, CSdef, and RBdef U2OS cell lines using the lncRNA-Seq service provided by Novogene. After quality control and rRNA depletion, directional libraries were prepared and sequenced on an Illumina NovaSeq X Plus platform using paired-end 150 bp reads. This yielded a total of approximately 247 million raw paired-end reads for WT (WT rep1: 78.8M pairs; WT rep2: 86.6M pairs; WT rep3: 81.6M pairs), 244 million raw paired-end reads for CSdef (CSdef rep1: 81.5M pairs; CSdef rep2: 81.4M pairs; CSdef rep3: 81.6M pairs), and 243 million raw paired-end reads for RBdef (RBdef rep1: 80.4M pairs; RBdef rep2: 82.7M pairs; RBdef rep3: 79.7M pairs). Demultiplexed FASTQ files were obtained and are publicly available as described below. Reads from the same sample sequenced across two flow cell lanes were merged prior to downstream analysis.

### Streamlined iCLIP (siCLIP)

A streamlined version of the individual-nucleotide resolution UV-crosslinking and immunoprecipitation (iCLIP) protocol was employed, adapted from^82^. U2OS cells expressing eGFP-tagged WT, CSdef, or RBdef FUS, alongside non-edited control (NEC) cells, were grown to ∼90% confluence and UV crosslinked on ice at 254 nm (150 mJ/cm²). WT eGFP-FUS cells without UV crosslink were included as control. Cells were lysed in lysis buffer containing protease inhibitor (Merck, #539134) and Turbo DNase (Ambion, #AM2238), and RNA was partially digested with RNase I (Ambion, #AM2295). Following centrifugation, cleared lysates were incubated with GFP-Trap M-270 magnetic beads (Proteintech, #gtd-20) to immunoprecipitate protein-RNA complexes. Beads were subjected to stringent washes and then treated with T4 PNK (New England Biolabs, #M0201) at pH 6.5 to dephosphorylate RNA 3′ ends. A Cy5-lableled barcoded adapter (Oligo ID: CS1) was ligated overnight at 16 °C using T4 RNA ligase (New England Biolabs, #M0437). Unligated adapters were removed by RecJ exonuclease (New England Biolabs, #M0264) treatment, and the complexes were eluted from beads in loading buffer, resolved by SDS-PAGE, and transferred to nitrocellulose membranes. Crosslinked complexes were visualised with a near-infrared imager (Odyssey Fc, LiCor) and excised from membranes. Membrane fragments were digested with proteinase K (Roche, #03115887001), and RNA was extracted using phenol-chloroform in Phase Lock Gel tubes (QuantaBio, #2302830) to ensure clean phase separation. Purification was performed using the Oligo Clean & Concentrator kit (Zymo, #D4060), which efficiently recovers small RNA species and RNA:DNA hybrids. Reverse transcription was carried out with AffinityScript reverse transcriptase (Agilent, #600107) using a biotinylated RT primer (Oligo ID: CS2), and excess primer was removed via hybridisation of a complementary oligonucleotide (Oligo ID: CS3) followed by exonuclease III (New England Biolabs, #M0206) digestion. The resulting biotinylated cDNA was captured on MyOne streptavidin beads (Invitrogen, #65001), ligated to a 3’ adapter (Oligo ID: CS4), and eluted by heating in nuclease-free water. Libraries were prepared by PCR amplification with indexed Illumina P5/P7 primers (18 cycles for WT, 19 cycles for CSdef and RBdef) and cleaned up using select-a-size DNA clean and concentrator columns (Zymo, #D4080). Library quality and concentration were assessed using a Bioanalyzer (Agilent), and sequencing was performed on an Illumina NextSeq 2000 platform in paired-end mode (Read 1: 75 cycles; Read 2: 40 cycles). This yielded a total of approximately 129 million raw paired-end reads for WT (WT rep1: 39.9M pairs; WT rep2: 46.9M pairs; WT rep3: 42.5M pairs), 111 million raw paired-end reads for CSdef (CSdef rep1: 43.3M pairs; CSdef rep2: 34.3M pairs; CSdef rep3: 33.0M pairs), and 147 million raw paired-end reads for RBdef (RBdef rep1: 48.1M pairs; RBdef rep2: 47.7M pairs; RBdef rep3: 50.9M pairs). Demultiplexed FASTQ files were obtained and are publicly available as described below.

### Differential expression analysis

After adapters trimming using TrimGalore (v0.6.6) and Cutadapt (v4.6), the reads were mapped to the human reference genome (GRCh38) using HISAT2 (v2.2.1) with the --dta option enabled to facilitate downstream transcript assembly. Transcript abundance was then quantified using StringTie (v1.3.5) guided by GENCODE version 33 gene annotations. GTF files produced by StringTie were processed using the prepDE.py script (provided by the StringTie developers at ccb.jhu.edu) to create a count matrix, and RPKM values were calculated using EdgeR (v4.6.2). Lowly expressed genes were filtered out by retaining only those with at least 1,000 total counts across all nine samples. Differential expression analysis was performed using DESeq2 (v1.48.1) and fold changes were moderated using DESeq2’s lfcShrink function with the “apeglm” method to improve effect size estimates for genes with high dispersion. Significant differential expression was defined by an absolute fold change > 2, adjusted p-value < 1E-05. Heatmaps were generated from variance stabilised transformed counts of selected genes and expression values were normalised by subtracting the gene-wise mean and dividing by the standard deviation across samples. A table listing log_2_ fold changes and adjusted p-values of all quantified genes is provided in Table S1.

### Gene ontology enrichment analysis

Gene Ontology (GO) enrichment analysis was performed using clusterProfiler (v4.16.0). Enrichment for Biological Process and Cellular Component terms was assessed using enrichGO with the Benjamini-Hochberg method for multiple testing correction.

### Quantification of exonic and intronic reads

To quantify gene expression using exonic and intronic reads only, custom GTF annotation files were prepared for exons and introns separately. Starting from the hg38 reference GTF file, exons were isolated, overlapping exons per gene merged using BEDTools (v2.31.1) and intronic regions defined as the genomic intervals between consecutive merged exons of the same gene. Aligned RNA sequencing reads were then quantified in exonic and intronic regions using featureCounts (Subread package, v.2.1.1) in paired-end, reverse-stranded mode, and summarised at the gene level. For each gene, RPKM values were then computed based on exonic or intronic counts. The RPKM values were subsequently log-transformed, and Z-scores were computed across conditions for visualisation as heatmap.

### Alternative splicing analysis

Alternative splicing was detected and quantified using rMATS (v4.3.0) with the --novelSS option enabled to detect novel splice sites, focusing on junction-spanning reads. Events with at least 10 inclusion and 10 skipping reads per sample were retained. Significant splicing changes were defined by an absolute ΔPSI > 10% and FDR < 0.01. A table listing ΔPSI values and false discovery rates of all quantified splicing events is provided in Tables S2-S5.

### Identification of UV crosslink sites

To emulate the iCLIP read structure, the barcode and UMI from read 2 were appended to the 5’-end of read 1. Restructured reads were then processed using the Flow.bio webserver: After adapter trimming using TrimGalore (v0.6.7) and Cutadapt (v3.4), the reads were first mapped to a small RNA index comprising rRNA and tRNA sequences using Bowtie (v1.3.0). Unmapped reads were subsequently mapped to the human reference genome (GRCh38) using STAR (v2.7.9a) with gene annotations from Ensembl release 109. Aligned reads were deduplicated based on unique molecular identifiers (UMIs) and their start positions. FUS crosslink sites were defined as the nucleotide position immediately upstream (-1) of each deduplicated read using BEDTools (v2.31.1). Given that the identified FUS crosslink sites were highly correlated between biological replicates, they were combined for further analysis. This resulted in a total of 18,284,974 unique crosslink sites for WT FUS, 7,776,767 for CSdef FUS and 6,203,397 for RBdef FUS. Downstream analysis of siCLIP data was performed as detailed below.

### Peak calling

FUS binding sites were identified using the Clippy (v1.5.0) peak-calling tool, applying minimum cDNA count thresholds of 5 for medium-confidence peaks. High-confidence peaks (referred to as ‘binding sites’) were defined with the following parameters: minimum cDNA count = 20, width = 0.7, rolling window size = 20, minimum prominence threshold = 3, minimum height threshold = 4. This resulted in 445,953, 259,587, and 214,460 medium-confidence peaks, along with 32,184, 13,903, and 6,241 high-confidence binding sites for WT, CSdef, and RBdef FUS.

### Motif discovery using positionally enriched k-mer analysis (PEKA)

To detect 5-mers that are enriched around crosslink clusters, we applied PEKA using default parameters to medium-confidence peaks. We restricted the analysis to introns, which harbour ∼80% of crosslinks. PEKA accounts for intrinsic crosslink biases by distinguishing thresholded crosslink sites within peaks - which represent strong, specific binding - from crosslink sites outside peaks, which indicate transient, non-specific binding and are used as background reference. After ranking all 5-mers by their PEKA scores for WT FUS, G- and C-rich motifs from the top 50 were selected and clustered for visualization as heatmap showing their relative occurrence around thresholded crosslink sites.

### FUS occupancy and differential binding at transcripts and binding sites

To assess FUS occupancy at the mRNA level, we quantified the number of unique crosslinks per protein-coding gene, normalised to library size. Since occupancy is influenced by mRNA expression levels, we restricted the analysis to genes that were not differentially expressed, defined by a fold change of less than 1.5 and an adjusted p-value greater than 0.001 in the DESeq2 analysis. To quantify differential binding of FUS mutants, we used the log_2_ transformed ratio of normalised occupancies (Mutant / WT). This identified 1,543 and 1,473 mRNAs with at least 2-fold increased or decreased binding for CSdef and RBdef FUS, respectively.

To assess FUS occupancy at individual binding sites, we first quantified the number of unique crosslinks for WT and each mutant FUS construct in the set of combined high-confidence FUS peaks (WT and CSdef, WT and RBdef). To account for differences in expression levels, we normalised the crosslink events within high-confidence peaks to the total crosslink count of the respective gene. Peaks that overlapped more than one annotated gene were discarded. Log-transformed ratios of normalised occupancies (Mutant / WT) were used to assess differential binding. This identified 12,789 and 10,560 binding sites with at least 2-fold increased or decreased binding for CSdef and RBdef FUS, respectively. Control sets of 24,962 and 23,085 unchanged binding sites (fold change smaller than 2) for CSdef and RBdef were defined for comparative analyses.

### Motif frequency analysis

To examine motif enrichment across distinct classes of FUS binding sites, we extended each identified peak by 5 nucleotides on both sides using BEDTools (v2.31.1) to capture motifs that may overlap peak boundaries. Corresponding nucleotide sequences were extracted from the human reference genome (GRCh38) using the BEDTools getfasta function with strand-specific output enabled. We then quantified 5-mer occurrences in each class of binding site using Jellyfish (v2.3.0). The resulting k-mer count tables from all six conditions (CS-sensitive, CS-induced, CS-insensitive, RB-sensitive, RB-induced, and RB-insensitive) were subsequently merged by k-mer identity. To enable direct comparison across categories, frequencies were normalised by dividing each k-mer count by the total number of counts per category. Normalised frequencies were used to compute per-category Z-scores, by subtracting the mean frequency across k-mers and dividing by the standard deviation. Z-scores thus represent motif enrichment relative to the global distribution of k-mer frequencies per category. 5-mers were subsequently categorised based on nucleotide composition (G-rich, C-rich, or other) and visualised using ggplot2 (tidyverse, v2.0.0).

### Secondary structure prediction

To investigate the secondary structure of the different class of FUS binding sites, peaks were first centred and extended by ±50 nucleotides around the midpoint. Nucleotide sequences were then extracted from the human reference genome (GRCh38) using BEDTools (v2.31.1) getfasta with strand specificity. RNA secondary structures were predicted using RNAfold (ViennaRNA package, v2.6.0), producing dot-bracket strings for each sequence. From these strings, we then computed per-nucleotide pairing probabilities by counting paired bases (“(” or “)”) for each position in the 101-nt window across all sequences and then normalising the frequencies by the total number of sequences. This generated per-position pairing probability profiles for each class of binding sites that were subsequently used for visualisation.

### RNA maps

To assess the positional enrichment of WT eGFP-FUS siCLIP binding sites in and around regulated exons, genomic occupancy profiles (RNA maps) were generated using R (v4.5.0). siCLIP peaks were imported from BED files produced by Clippy (v1.5.0, parameters: mb = 10, w = 0.7, n = 20) using the rtracklayer package (v1.68.0) and converted into a GRanges object from GenomicRanges (v1.60.0). Cassette exon coordinates were obtained from the differential splicing analysis using rMATS. Exons that were over 20 nt in length and not significantly regulated in any condition (CSdef or RBdef) were selected to define a background set of unaffected exons. For each exon, strand-aware 300-nt intronic and 20-nt exonic windows were generated at both splice junctions using the promoters() and flank() functions. The regions surrounding each regulated exon were divided into 10-nt bins with tile(), producing a uniform set of windows spanning exon-intron boundaries. siCLIP peaks overlapping each 10-nt bin were then identified using findOverlaps() with strand specificity enforced. A binary indicator (1 = overlap, 0 = no overlap) was assigned to each bin to capture the presence of binding sites. Normalised peak density at each position was quantified by summing the number of bins containing at least one peak and normalising by the total number of exons in the set. The resulting peak density profiles were visualised using ggplot2 (tidyverse, v2.0.0), applying locally weighted smoothing (span = 0.2) to display normalised peak coverage along regulated exons.

